# Reprogramming of stromal fibroblasts by chemotherapy-induced secretion of IFNβ1 drives re-growth of breast cancer cells after treatment

**DOI:** 10.1101/2020.08.05.238436

**Authors:** Ana Maia, Zuguang Gu, André Koch, Rainer Will, Matthias Schlesner, Stefan Wiemann

**Affiliations:** Division of Molecular Genome Analysis, German Cancer Research Center (DKFZ), Im Neuenheimer Feld 580, 69120 Heidelberg, Germany; Faculty of Biosciences, University of Heidelberg, Germany; Computational Oncology, Molecular Diagnostics Program, National Center for Tumour Diseases (NCT) and German Cancer Research Center (DKFZ), Im Neuenheimer Feld 280, 69120 Heidelberg, Germany; DKFZ-HIPO (Heidelberg Center for Personalized Oncology), 69120 Heidelberg, Germany; Department of Women’s Health Tübingen, Eberhard-Karls-University Tübingen, 72076 Tübingen, Germany; Genomics and Proteomics Core Facility, German Cancer Research Center (DKFZ), Im Neuenheimer Feld 580, 69120 Heidelberg, Germany; Bioinformatics and Omics Data, German Cancer Research Center (DKFZ), 69120 Heidelberg, Germany

**Keywords:** Breast cancer, fibroblasts, tumour microenvironment, chemotherapy, IFNβ1

## Abstract

Chemotherapy is still the standard of care for a large number of aggressive tumours including breast cancer. In breast cancer, chemotherapeutic regimens are administered in intervaled cycles of the maximum tolerated dose, allowing cancer cells to re-grow or adapt during the resting periods between cycles. However, how stromal fibroblasts impact the fate of cancer cells after chemotherapy treatment remains poorly understood. We show that cancer cells utilize paracrine signalling with stromal fibroblasts to drive their recovery after treatment withdrawal. Secretion of IFNβ1 by cancer cells after treatment with high doses of chemotherapy instigates the acquisition of an anti-viral state in stromal fibroblasts associated with the expression of several interferon stimulated genes (ISGs), including numerous pro-inflammatory cytokines. This crosstalk is an important driver of the expansion of breast cancer cells after chemotherapy and blocking of IFNβ1 in tumour cells abrogated their increased recovery potential. Analysis of human breast carcinomas supports the proposed role of IFNβ1 since its expression is inversely correlated with recurrence free survival (RFS). Moreover, expression of the interferon signature identified in stromal fibroblasts is equally associated with higher recurrence rates and a worse outcome in breast cancer patients. Our study unravels a novel paracrine communication between cancer cells and fibroblasts that ultimately results in the escape of malignant cells to treatment. Targeting of this axis could potentially improve the outcome of breast cancer patients to chemotherapy treatment.

## Introduction

Breast cancer is still the most common type of malignancy and is responsible for the highest number of cancer-related deaths among woman^1,2^. For a large number of cancer patients, chemotherapy is still the standard of care and despite the initial good response rates, patients often return to the clinic with relapses. Therapeutic failure often arises from the expansion of clones of resistant cancer cells. These cells can be present in the initial tumour mass, but they can also arise later through adaption. Nevertheless, non-resistant tumour cells that are not eliminated at the time of treatment also play an important role in the development of relapses. In fact, drug tolerant tumour cells that do not undergo apoptosis or senescence after treatment can persist in the patient body, adapt and later give rise to tumour recurrences^3^. In breast cancer, chemotherapy is still given as cycles of maximum tolerated doses, which allows cancer cells to re-grow and adapt during the resting periods. How cancer cells survive and escape therapy and what role stromal cells play in this process is still unclear.

The tumour microenvironment is an important player during all steps of tumour development and progression and in breast cancer the majority of the stroma is constituted by cancer-associated fibroblasts (CAFs). CAFs originate from a variety of cell types but the most prominent source are resident fibroblasts that become activated and then display an enhanced secretory profile^4^. By secreting a variety of growth factors, cytokines and extracellular matrix, CAFs are known to exert their control over an array of processes, including resistance to chemotherapy^5,6^. On the other hand, the phenotype and activation profile of CAFs are equally influenced by their surroundings, especially, but not exclusively, by cancer cells^4^. In the context of chemotherapy, it is increasingly evident that these agents strongly modulate the secretory profile of cancer cells^7^. Alterations in the array of secreted factors dictates the interaction between cancer and stromal cells. Understanding how these cell types communicate in the context of chemotherapy is essential to understand therapeutic outcome and may provide insights into promising new drug targets.

Type I interferons (IFN) belong to the cytokine family and are secreted signalling molecules that are involved in the response of the immune system to infections^8^. Recognition of pathogen-associated molecular patterns (PAMPs) by specialised receptors named pattern recognition receptors (PRRs), leads to the expression of type I IFN^9,10^. It is also known that this signalling is not only activated by foreign organisms, but that PRRs can also recognise self-derived danger signals named damage-associated molecular patterns (DAMPs)^11^. Such danger molecules often derive from damaged or dying cells making tumours often DAMP-rich environments. Cytoplasmic nuclei acid sensors that recognise these danger molecules and induce type I IFN expression have been broadly described to be strongly activated in tumours as a response to ionizing therapy^12–15^. All type I IFNs, which comprise 13 IFNa isoforms and IFNβ1, bind a common receptor formed by the heterodimer of IFNAR1 and IFNAR2. Binding of type I IFNs to its receptor triggers a signaling cascade that culminates with the activation of signal transducer and activator of transcription (STAT), which then mediates the transcription of interferon-stimulated genes (ISGs)^16,17^. ISGs expression in cancer has been associated with a gene signature that predicts response to both radiation and chemotherapy^18,19^. Expression of these ISGs is often driven by an anti-viral response in cancer cells and was later shown to be induced in a paracrine manner by fibroblasts in cancer cells to promote chemoresistance^5^. The role of IFN signalling in cancer and its impact in dictating response to chemotherapy is still poorly understood with studies showing both tumour-inhibitory^20–22^ and -promoting^23–25^ activities.

In this study, we set out to investigate the impact of stromal fibroblasts on the fate of cancer cells after exposure to high doses of cytotoxic drugs and unravel the communication axis between these two cell types in a chemotherapy-dependent context. We show that stromal fibroblasts strongly increase the re-growth of cancer cells after treatment with high doses of drugs and identified IFNβ1 as an important factor of the therapy-induced tumour cell secretome. Secretion of IFNβ1 by chemotherapy-treated cancer cells led to an anti-viral response in stromal fibroblasts, associated with the expression of several ISGs such as *DDX58, IFIH1, ISG15* and *OAS1.* Ablation of cancer cell-derived IFNβ1 resulted in the abrogation of this activation state in fibroblasts and more importantly decreased the recovery potential of cancer cells after chemotherapy treatment. Thus, our study defined a novel pro-tumorigenic state of fibroblasts that is dependent on type I IFN signalling and that drives the escape of cancer cells from therapy.

## Results

### Fibroblasts promote the recovery of cancer cells after high doses of chemotherapy

Fibroblasts are important components of the tumour stroma, especially in breast cancer where they are the most prominent cell type surrounding neoplastic cells. We sought to investigate if stromal fibroblasts are capable of affecting the fate of cancer cells after exposure to chemotherapeutic agents. For this, we used primary fibroblasts isolated from breast cancer patients from either tumour sites (CAF) or non-malignant sites (NF). The fibroblast identity of the primary cells used was validated, and a representative image is shown in Fig S1A. Additionally, levels of the commonly used cancer-associated fibroblast marker, aSMA, were also investigate (Fig S1B). Initially, the response curves of the breast cancer cell lines MCF7, HS578T and SKBR3 to commonly used chemotherapy drugs – epirubicin and paclitaxel – were determined (Fig S1C-D). Cancer cells exposed to high chemotherapy doses were allowed to recover either in mono-culture (MC) or indirect co-culture (CC) with primary fibroblasts isolated from breast cancer samples as shown in Fig 1A. Briefly, cancer cells were treated with IC90 doses of chemotherapy for three days after which the drug was removed, and they were either cultured in the presence or absence of fibroblasts. Treatment of MCF7, HS578T and SKBR3 with chemotherapy resulted in the recovery of a very small fraction of cells (bellow 0.1% of total seeded cells). Analysis of the number and size of the cancer cells colonies revealed that co-culture with all primary fibroblast lines tested significantly increased the number of cells that recovered after paclitaxel treatment (Fig 1B-E; Fig S1E). This was independent if fibroblasts had been isolated from a non-malignant or a tumour site (NF or CAF, respectively). A similar effect in the recovery of cancer cells treated with epirubicin was observed (Fig S1F-G). On the other hand, co-culture of untreated MCF7 with CAF1 did not increase the colony formation potential of cancer cells (Fig S1H).

**Figure 1.**
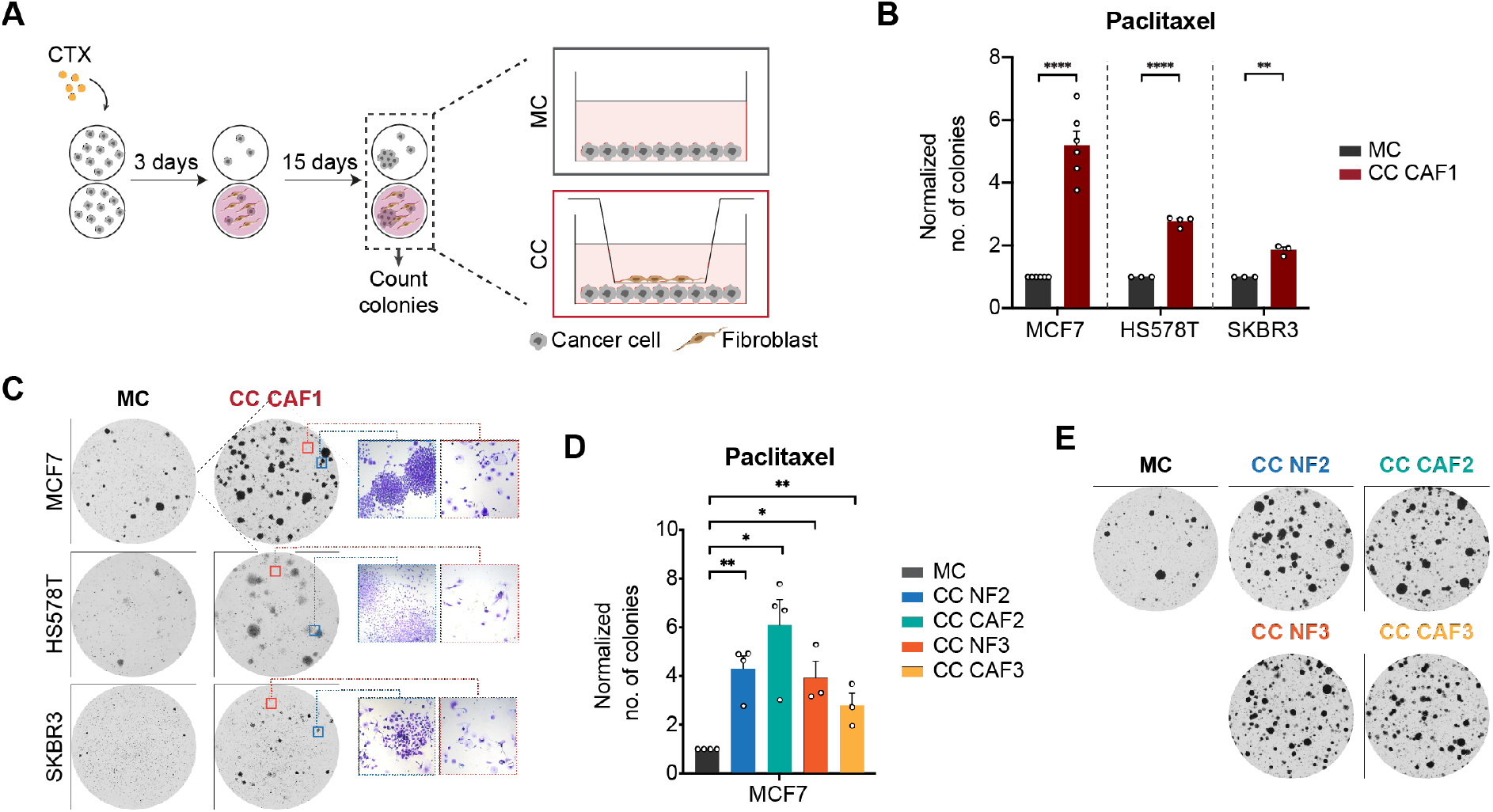
Fibroblasts promote the recovery of cancer cells after high-dose chemotherapy. **A**: Schematic overview of recovery assay. Cancer cells were treated for three days with IC90 doses of chemotherapy (CTX) and allowed to recover for two weeks in the absence (MC) or presence (CC) of fibroblasts. Fibroblasts were seeded in trans-wells and communicated with cancer cells via secreted factors as depicted in scheme. **B-C**: Quantification of breast cancer cell lines colonies in MC and CC with CAF1 at the end of recovery after treatment with IC90 doses of paclitaxel (**B**). Each dot represents an independent biological replicate (MCF7: n= 6, HS578T: n=4, SKBR3 n=3). Representative pictures of wells used for colony quantification are shown in **C**. Crystal violet staining under microscope (10x magnification) showing an area containing viable, proliferating colonies and arrested cells are shown in the blue and orange squares, respectively. **D-E**: Recovery assay of MCF7 exposed to 4 nM paclitaxel in MC or CC with CAF2/NF2 and CAF3/NF3. Quantification of the number of colonies at the end-point is shown in **D** and representative pictures of the wells in **E**. Four independent biological replicates (n=4) were done for fibroblast pair #2 and three independent replicates (n=3) for fibroblast pair #3. Data is shown as mean ± SEM. P values were calculated using unpaired two-tailed t-test on biological replicates. * p<0.05, ** p<0.01, **** p<0.0001. Each dot represents an independent replicate.

To determine if the cells that recovered after chemotherapy treatment were resistant to the treatment and if co-culture with fibroblasts affected their response to the treatment, colonies from both mono-culture (Col-MC) and co-culture (Col-CC) that survived paclitaxel-treatment were expanded (Fig S2A). Drug response assays revealed that colonies derived from MC and CC responded in a very similar fashion to paclitaxel treatment not only between each other but also when compared to the parental cell line (Fig S2B-C), suggesting that recovery after treatment was not associated with resistance.

Combined these results indicate that the presence of stromal fibroblasts promotes the survival and re-growth of non-resistant cancer cells after chemotherapy treatment. To investigate the underlying mechanisms, additional phenotypic assays and RNA-sequencing of both cancer cells and fibroblasts were performed.

### Fibroblasts drive cell cycle re-entry of cancer cells after chemotherapy

Fibroblasts can modulate different aspects of tumour progression. Several studies have shown that the presence of fibroblasts can prevent cancer cells from undergoing programmed cell death after chemotherapy treatment^5,26^. To investigate if co-culture with fibroblasts modulated the levels of apoptosis of cancer cells during the recovery period, the number of apoptotic cells was measured at different time points. DAPI labelling was used as a readout of cell death and the DAPI-negative fraction was quantified by flow cytometry (Fig S3A). No difference in the percentage of alive cancer cells between mono-culture and co-culture was observed in any of the cell lines tested (Fig 2A and Fig S3B), indicating that fibroblasts are not protective against apoptosis. To further explore the role of fibroblasts during recovery, cell cycle profiling of cancer cells was done. EdU labelling revealed an increase in the percentage of cancer cells going through S-phase in the presence of fibroblasts. This effect was observed at several time points of recovery (Fig 2B and Fig S3C-D). In addition, the number of senescent MCF7 cells was reduced in co-culture conditions during the recovery period (Fig S3E). Finally, analysis of the different cell cycle phases using the FUCCI system^27^ also supported these findings. Very briefly, cells with red nuclei (Cdt1-RFP) were counted as G1, while cells with green nuclei (Geminin-GFP) were counted as S-G2-M. After eight days of co-culture with CAF1 there was a reduction in the percentage of cells expressing Cdt1 which was accompanied by an increase in the fraction of GFP-positive nuclei, pointing to a decrease in the number of G1 arrested cells (Fig S3F-H). These data indicate that fibroblasts promote the cell cycle re-entry of cancer cells after treatment with high doses of chemotherapy.

**Figure 2.**
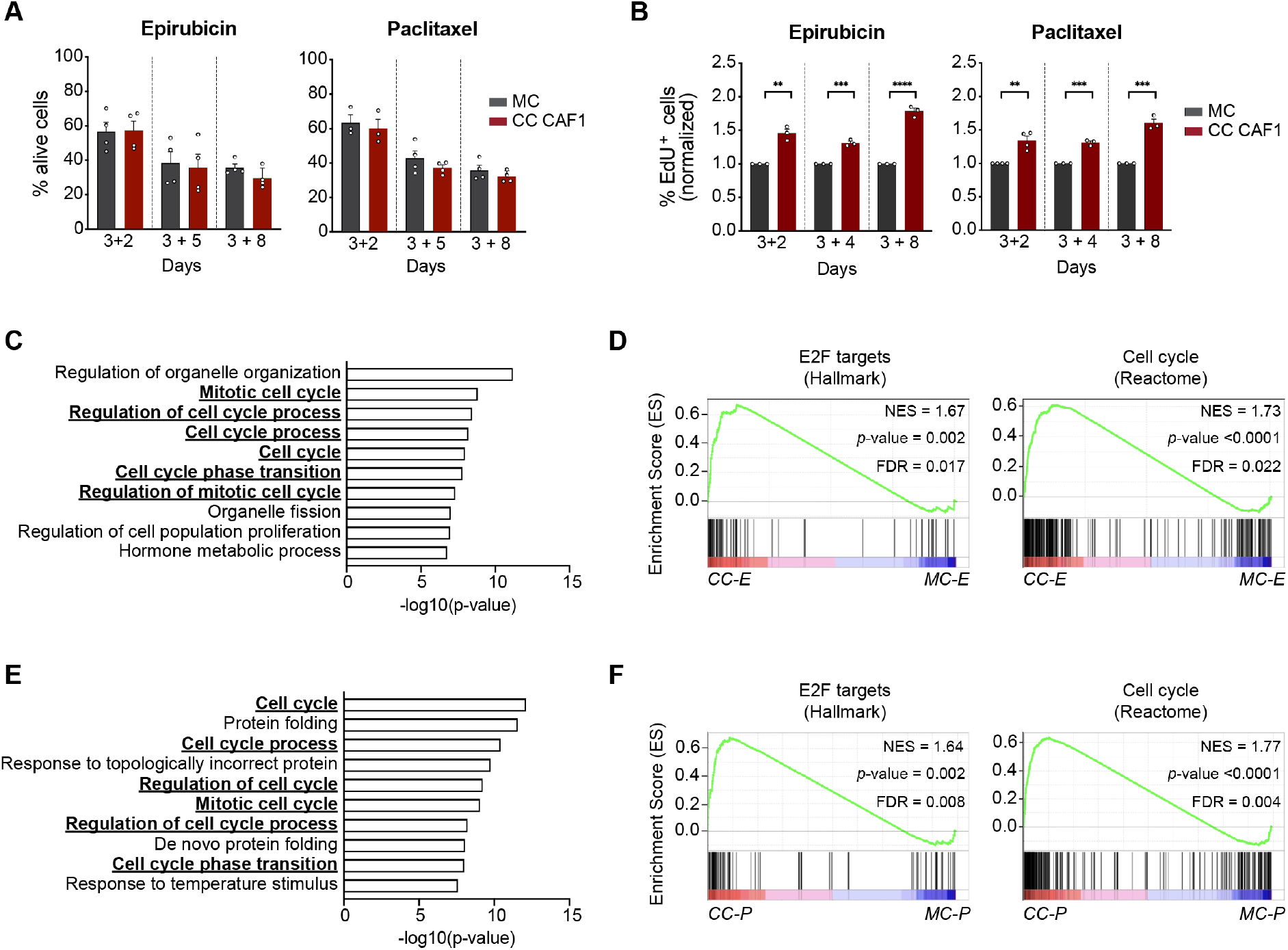
Fibroblasts promote the re-entering in cell cycle of cancer cells after chemotherapy treatment. **A**: Apoptosis measurement of MCF7 in mono-culture (MC) and co-culture with CAF1 (CC CAF1). Single cells negative for DAPI staining were counted and quantified by flow cytometry. Graphs show % of cells alive after treatment with either epirubicin (left) or paclitaxel (right) at different time points of recovery – after two (3+2), five (3+5) or eight (3+8) days of drug withdrawal. **B**: Quantification of EdU-labelled nuclei in MCF7 cells. % of EdU-positive cells was calculated by dividing labelled cells by total cells (DAPI staining). Values were normalized to monoculture condition and are shown in the graphs for cancer cells treated with epirubicin (left) and paclitaxel (right). In **A** and **B**, data is shown as mean ± SEM, p values were calculated using unpaired two-tailed t-test in biological replicates. ** p<0.01, *** p<0.001, **** p<0.0001. Each dot represents an independent replicate (n≥3). **C**: Gene ontology (GO) analysis of upregulated genes in epirubicin-treated MCF7 in co-culture with CAF1 (compared to MCF7 in mono-culture) for function (GO: Biological process) – top 10 list. Terms related to cell cycle are underlined and highlighted in bold. **D**: Enrichment of proliferation signatures (E2F targets – HALLMARK, and Cell cycle – REACTOME) in epirubicin-treated MCF7 in CC (CC-E) with CAF1 compared to MCF7 in mono-culture (MC-E). NES = normalized enrichment score, FDR = false discovery rate. P values were determined by random permutation tests. **E**: Gene ontology (BP) analysis of upregulated genes in paclitaxel-treated MCF7 in co-culture with CAF1 (compared to MCF7 in MC) – top 10 list. Terms related to cell cycle are underlined and highlighted in bold. **F**: Enrichment of proliferation signatures E2F targets – HALLMARK, and Cell cycle – REACTOME) in paclitaxel-treated MCF7 in CC with CAF1 compared to MCF7 in MC. NES = normalized enrichment score, FDR = false discovery rate. P values were determined by random permutation tests.

To further explore the crosstalk between cancer cells and fibroblasts in this context, we performed transcriptomic analysis by RNA-sequencing of both MCF7 and CAF1 (see Fig S4A for experimental layout). Treatment of MCF7 with epirubicin and paclitaxel led to the upregulation (fold change >2, p<0.05) of 2513 and 1533 genes, and downregulation (fold change <0.5, p<0.05) of 1014 and 338 genes, respectively (Fig S4B). The vast majority of deregulated genes was shared by both treatments, pointing to the acquisition of a common transcriptional profile by cancer cells after exposure to the two cytotoxic agents. Several genes involved in DNA damage sensing pathways, including the p53 pathway, were upregulated in treated conditions compared to untreated while downregulated genes included genes involved in cell cycle progression (e.g., mitotic genes such as *BUB1* and *TTK),* confirming the induction of a proliferation arrest in cells by both treatments (Fig S4C). Gene ontology (GO) term enrichment analysis of the transcriptome of chemotherapy-treated MCF7 in mono-culture versus co-culture revealed that the presence of fibroblasts induced a signature of genes involved in cell cycle progression in cancer cells (Fig 2C and Fig 2E for epirubicin and paclitaxel treatment, respectively), confirming the results from the phenotypic assays (Fig 2B). This signature was not observed in untreated cancer cells that had been in co-culture with fibroblasts (Fig S4D). In addition, gene set enrichment analysis (GSEA) showed that only in chemotherapy-treated MCF7 in co-culture with CAF1, but not untreated MCF7, there was an enrichment in proliferation signatures (Fig 2D and Fig 2F for epirubicin and paclitaxel treatment, respectively. See <u>Supplementary Table 1</u> for list of top 10 HALLMARKS and REACTOME in three conditions (epirubicin – E, paclitaxel – P, untreated – U) with FDR <0.1).

The above findings suggest a model in which stromal fibroblasts promote the cell cycle re-entry of cancer cells, likely by providing them with factors that allow cancer cell survival and proliferation after treatment with chemotherapeutic agents.

### Chemotherapy-treated cancer cells induce an anti-viral state in fibroblasts

Gene ontology (GO) analysis for both function (GO: Biological process) and localisation (GO: Cellular component) of the top 2000 upregulated genes in paclitaxel- and epirubicin-treated MCF7 showed that these gene sets were enriched for secreted factors (Fig 3A and Fig S5A), which points to a reprogramming of the secretory profile of cancer cells after chemotherapy treatment. To understand how the chemotherapy-induced secretome of cancer cells affected the fibroblasts that were expose to it, we next explored the transcriptomic profile of fibroblasts in co-culture with untreated and treated cancer cells. Comparison between these conditions revealed a strong induction of an anti-viral response in fibroblasts in co-culture with cancer cells treated with both paclitaxel and epirubicin (Fig 3B and Fig S5B, respectively). This antiviral state in fibroblasts was accompanied by the acquisition of an inflammatory response signature (Fig 3C and Fig S5C-D). Moreover, we investigated which transcription factors and motifs might be involved in the regulation of the upregulated genes in the fibroblasts in coculture with chemotherapy-treated cancer cells and identified interferon regulatory factors (IRF) and interferon responsive sequence element (IRSE) motif as the top hits (Fig 3D and Fig S5E). These findings indicate that chemotherapy-treated cancer cells trigger the acquisition of an anti-viral state in fibroblasts characterized by the expression of numerous inflammatory modulators including several ISGs. To validate these results, expression of *DDX58, IFIH1, ISG15* and *OAS1* in fibroblasts was assessed by reverse transcription with quantitative PCR (RT-qPCR) and used as a readout of the anti-viral state (Fig 3E and Fig S5F). Strikingly, the expression of the genes in CAF1 was not induced or only weakly upregulated when fibroblasts were in co-culture with untreated cancer cells (Fig 3G-H). In contrast, treatment of cancer cells with paclitaxel prior to the co-culture led to a significant increase in the expression of all four anti-viral genes, validating the results above. It has been previously demonstrated that treatment of primary fibroblasts with high doses of chemotherapy can induce a pro-inflammatory state in fibroblasts^28^. Notably, in this context chemotherapy treatment of CAF1 did not induce the expression of any of the ISGs analyzed, except for *OAS1* upon paclitaxel-treatment (Fig S5G).

**Figure 3.**
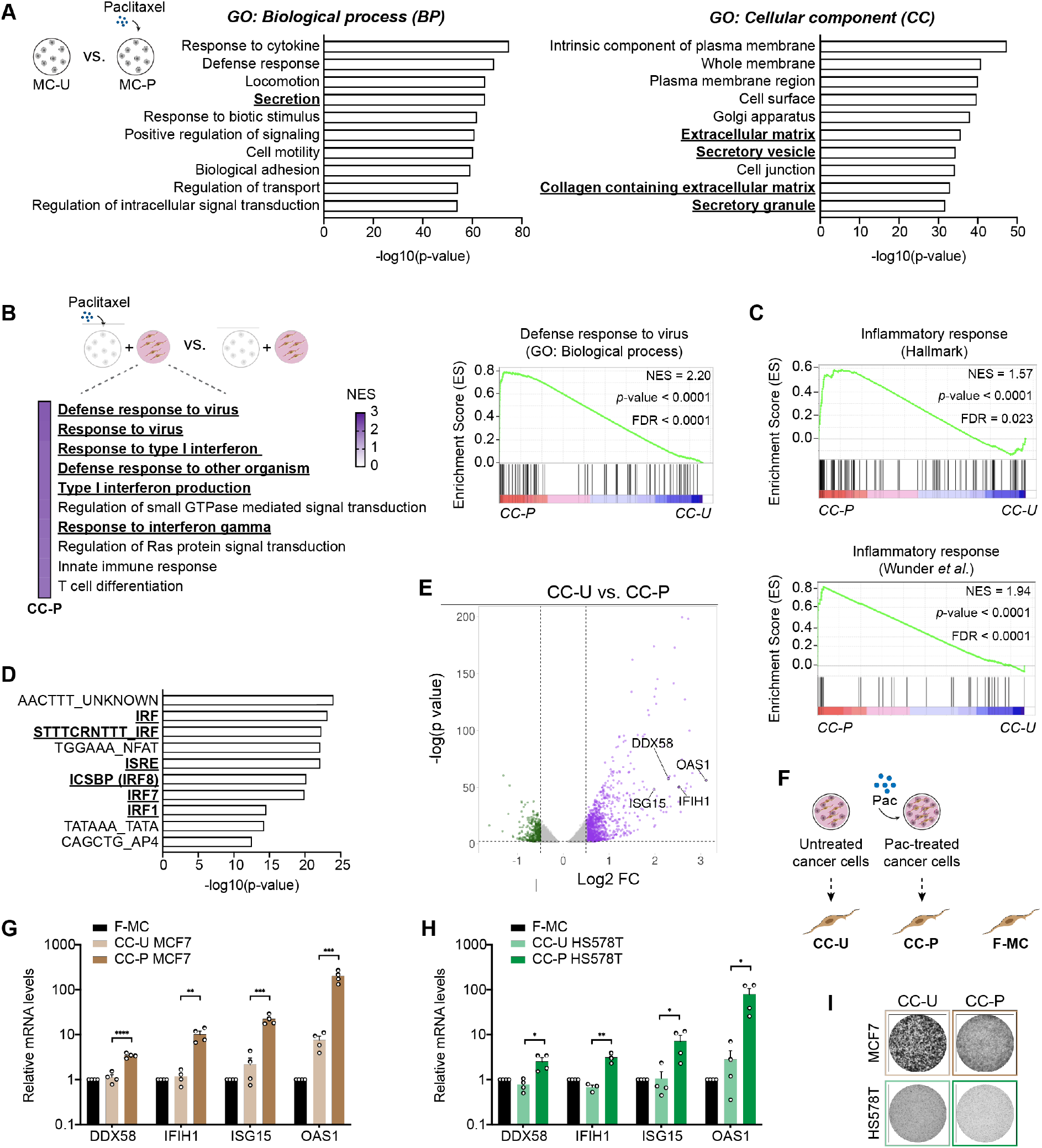
Fibroblasts acquire an anti-viral state after co-culture with chemotherapy-treated cancer cells. **A**: Top 10 enriched GO terms for biological process (BP) (left panel) and cellular component (CC) (right panel) of top 2000 upregulated genes in paclitaxel-treated MCF7 compared to untreated. Terms related to secreted factors are highlighted in bold. **B**: GSEA analysis of upregulated genes in CAF1 in CC-P compared to CC-U. Top 10 significantly enriched GO terms enriched are shown. NES = normalized enriched score. Terms related to anti-viral response are highlighted. Right panel shows enrichment for viral responses signature in CAF1 in CC-P versus CC-U. FDR = false discovery rate. P values were calculated by random permutation tests. **C**: Upregulated genes in CAF1 in co-culture with paclitaxel-treated cancer cells (CC-P) are involved in inflammatory response determined by enrichment analysis (HALLMARK and Wunder *et al.* ^53^). **D**: Top 10 over-represented transcription factors and motifs involved in the regulation of genes upregulated in CAF1 in co-culture with paclitaxel-treated cancer cells compared to CAF1 in co-culture with untreated cancer cell. Transcription factors known to be involved in anti-viral response are underlined and highlighted in bold. **E**: Volcano plots of differently expressed genes in CAF1 in CC-P compared to CC-U. Upregulated genes are shown in purple and downregulated genes in green. *DDX58, IFIH1, ISG15* and *OAS1* are highlighted. Volcano plot was designed using *VolcaNoseR^54^* tool. **F**: Schematic overview of experimental conditions. CAF1 were grown in mono-culture (F-MC), in co-culture with paclitaxel-treated cancer cells (CC-P) or in co-culture with untreated cancer cells (CC-U). **G-I**: Validation of anti-viral like state in CAF1 in co-culture (CC) with MCF7 (**G**) and HS578T (**H**) (n=4 for each cell line). Expression of *DDX58, IFIH1, ISG15* and *OAS1* in CAF1 was investigated by qRT-PCR and was normalised to F-MC. Representative pictures of cancer cells in untreated and paclitaxel conditions at the end-point of co-culture are shown in **I**. Data is shown as mean ± SEM, p values were calculated using unpaired two-tailed t-test in biological conditions. * p<0.05, ** p<0.01, *** p<0.001, **** p<0.0001. Each dot represents an independent replicate.

Taken together, we show that fibroblasts are reprogrammed by chemotherapy-treated cancer cells. Fibroblasts in this context acquire a similar transcriptional profile to cells exposed to viruses with increased expression of several pro-inflammatory modulators.

### Anti-viral state in fibroblasts is independent of nucleic acid sensing pathways

We next sought to determine the mechanism that drives the anti-viral state in fibroblasts. To understand if the transcriptional reprogramming of fibroblasts was induced directly by factors secreted by cancer cells after treatment with chemotherapy or if the crosstalk between cancer cells and fibroblasts played any role in promoting the expression of anti-viral genes, the supernatants from untreated or paclitaxel-treated cancer cells were collected and directly added to fibroblasts (Fig S6A). As in all previous experiments, the chemotherapeutic drugs had been removed after three days and were not present in the media at the time the supernatants were added to the fibroblasts. The exposure of CAF1 to the supernatant of both MCF7 and HS578T previously treated with IC90 doses of paclitaxel was sufficient to induce the expression of the anti-viral genes (Fig S6B). As before, this increase was absent in fibroblasts exposed to the conditioned-media of untreated cancer cells. Furthermore, upregulation of the anti-viral genes was observed in fibroblasts that had been isolated from both non-malignant and tumour tissue, indicating that these were equally reprogrammed by the secretome of paclitaxel-treated cancer cells (Fig S6C). This analysis supports the concept that chemotherapeutic drugs modify the secretome of cancer cells in breast cancer. We further demonstrate that this newly acquired TCS can then affect their communication with the microenvironment and highjack the fibroblasts that are exposed to it to induce the anti-viral state.

Previous studies show that the expression of anti-viral genes can result from the activation of multiple pathways. The two main nucleic acid sensing pathways are activated by RIG-I and Toll-like receptors, which directly bind and sense viral DNA or RNA. In the tumour context these pathways can be activated by exosomes^5^ or by DAMPs derived from cancer cells which can then be recognised by neighbouring cells^29^. To understand if these pathways were involved in the induction of the anti-viral like state in the fibroblasts, the supernatants from MCF7 and HS578T treated with paclitaxel were collected and subjected to different treatments (See Fig 4A for schematic layout). RNase and DNase treatment of the conditioned-media did not have any impact in the expression of anti-viral genes in the fibroblasts. By contrast, boiling the conditioned-media for five minutes completely abolished the up-regulation of the four anti-viral genes (Fig 4B). Additionally, the supernatant collected from paclitaxel-treated cancer cells was submitted to ultracentrifugation and the exosome-enriched fraction (Exo^+^) and exosome-depleted fraction (Exo^-^) were collected and added to fibroblasts. Only the exosome-depleted fraction was capable of inducing the expression of *DDX58, IFIH1, ISG15* and *OAS1* to a similar extent as the ‘full’ conditioned-media (Fig S6D). These results suggest that the anti-viral state is not induced by exosomes or DAMPs secreted by cancer cells after chemotherapy treatment but rather by a protein factor. To further confirm these observations, we took the complementary approach of silencing the nuclei acid sensing pathways in fibroblasts using RNA-interference (See Fig 4C for schematic layout). Knock-down (KD) of one or both RNA cytosolic receptors of the RIG-I pathway – *DDX58* and *IFIH1* – had no impact in the anti-viral like state (Fig 4D and Fig S6E for KD efficiency). Furthermore, silencing of *MYD88,* an important mediator of Toll-like receptor signalling, was also not enough to abolish the induction of the anti-viral genes in fibroblasts (Fig 4D and Fig S6E for siRNA silencing efficiency).

**Figure 4.**
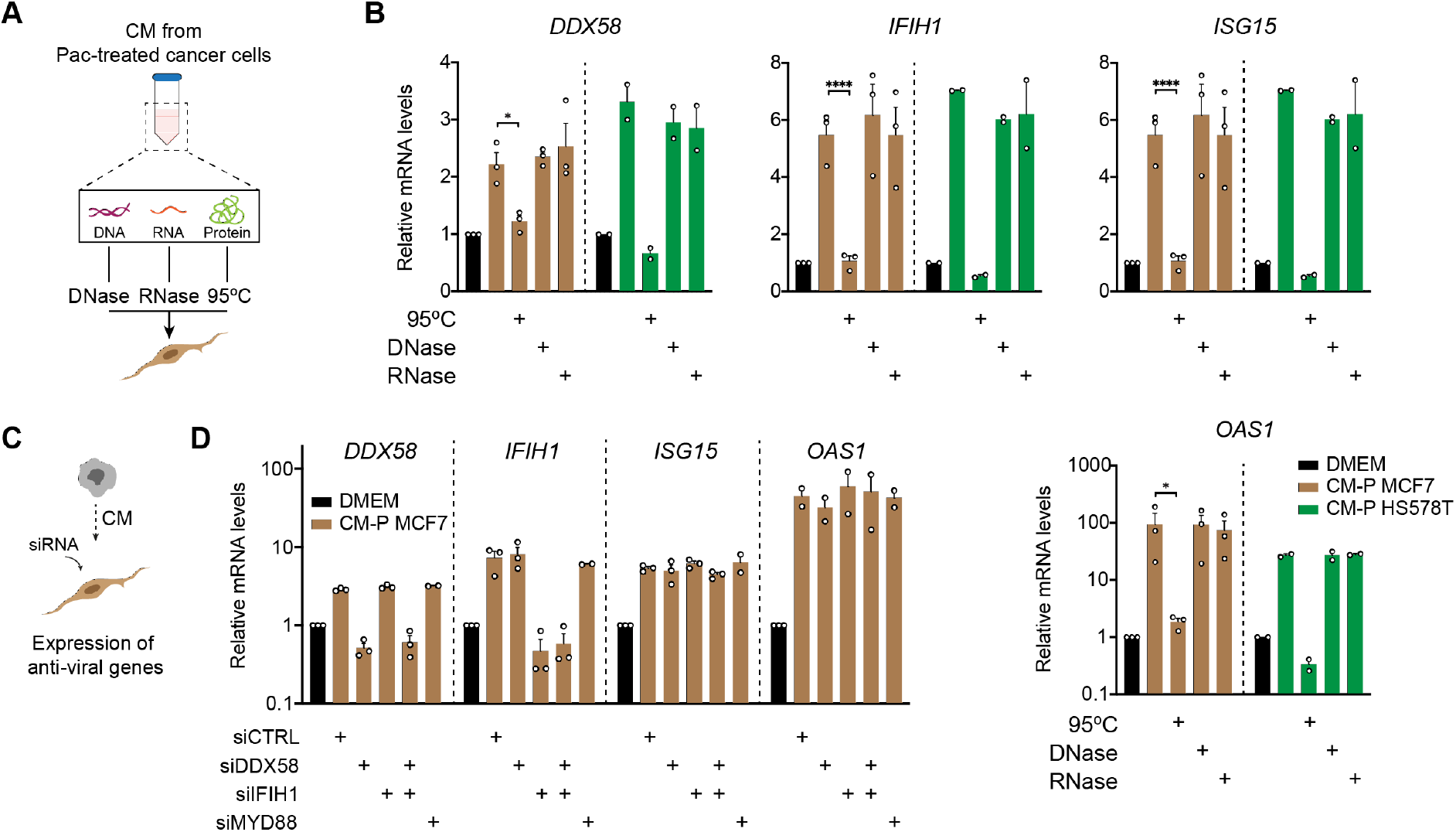
Anti-viral like state in fibroblasts is independent of nuclei acid sensing pathways. **A**: Schematics of the different treatments to which the supernatant of cancer cells was exposed. Collected conditioned-media (CM) from treated cancer cells was either exposed to DNase, RNase or 95°C treatment before addition to fibroblasts. **B**: RT-qPCR analysis of anti-viral genes in CAF1 after exposure to the conditioned-media without any treatment, after boiling (95°C), after DNASE or RNASE treatment. CM from both MCF7 (n=3) and HS578T (n=2) treated with 4 and 8 nM paclitaxel, respectively, was used. **C**: Layout of experimental setup for nuclei acid sensing receptors KD experiments. **D**: KD of nucleic acid sensing pathways in CAF1 and its impact in the expression of anti-viral genes measured by RT-qPCR. Each dot represents an independent biological replicate. Data is shown as mean ± SEM, p values for experiments with n>2 were calculated using unpaired two-tailed t-test in biological replicates. * p<0.05, **** p<0.0001.

Collectively these data indicate that fibroblasts are modulated by a soluble protein factor that is secreted by cancer cells after chemotherapy treatment rather than by nuclei acids or exosomes that activate nuclei acid sensing pathways.

### Chemotherapy-induced secretion of IFNβ1 by cancer cells induces anti-viral state in fibroblasts and promotes recovery

Based on the evidence that the anti-viral state in fibroblasts is not mediated by DNA or RNA-sensing pathways but by a protein, we continued to explore our transcriptomic data to identify candidate factors. A strong enrichment in interferon response signaling in fibroblasts in co-culture with chemotherapy-treated cancer cells was observed (Fig 5A and Fig S7A). Comparison between the expression of interferon transcripts in untreated MCF7 cells and their treated counterparts showed a strong upregulation of type I – *IFNB1* – and type III – *IFNL1, IFNL2, IFNL3* – interferons after treatment (Fig 5B). No reads were detected for the majority of other IFNs, including all 13 isoforms of *IFNA* and *IFNG.* Upregulation of *IFNB1* and *IFNL1* was validated by RT-qPCR in MCF7 and HS578T (Fig S7B). Both type I and type III interferons are potent inducers of the anti-viral state and are known to trigger the expression of ISGs. However, contrary to type I interferon which is known to mediate this state in any type of tissue due to the ubiquitous expression of its receptor – the IFNAR1-IFNAR2 heterodimer –, type III interferon activity seems to be compartmentalised to epithelial cells^30^. Type III interferon receptor is formed by the heterodimer of IL10RB and IL28RA. The latter receptor is reported to only be expressed by epithelial cells. Accordingly, no reads were detected in fibroblasts in our RNA-sequencing (<u>data not shown</u>). Nonetheless, we modulated the expression of both IFNβ1 and IFNλ1 in cancer cells in order to understand if they were responsible for the activation state observed in fibroblasts (Fig 5C). Blockade of IFNβ1 using RNA-interference or a blocking antibody, led to an almost complete abrogation of the upregulation of anti-viral genes in fibroblasts (Fig 5D-E and Fig S7E; Fig S7C-D for siRNA silencing efficiency), whereas silencing of IFNλ1 in cancer cells had no effect (Fig 5D-E; Fig S7C-D for siRNA silencing efficiency). After binding to its receptor, type I interferon induces a cascade of events via JAK proteins that culminates in the phosphorylation of the transcription factor STAT1, which then translocates to the nucleus where it induces the expression of interferon target genes, including *STAT1* itself. We saw a strong upregulation in the expression of STAT 1 as well as its activation marker – serine 727 phosphorylation (S727) in CAF1 in co-culture with paclitaxel-treated MCF7 (Fig 5F). Consistent with these results, depletion of the IFNβ1 receptor – IFNAR1 – in fibroblasts, similarly abrogated the anti-viral state observed (Fig S7G-H and Fig S7F for siRNA silencing efficiency). This was further confirmed with two independent siRNAs targeting siIFNAR1 (Fig S7J-K). Moreover, the increase in expression of STAT1 in CAF1 mediated by co-culture with paclitaxel-treated MCF7 was abrogated upon silencing of *IFNAR1* in the fibroblasts (Fig S7I). To test whether this communication axis played a role in the recovery of cancer cells after chemotherapy treatment, we then silenced IFNβ1 in cancer cells and investigated its impact on the colony formation potential of these cells after paclitaxel treatment. Depletion of IFNβ1 was sufficient to decrease the recovery capacity of cancer cells in the presence of fibroblasts (Fig 5G-H), indicating that this paracrine communication between cancer cells and stromal fibroblasts plays an important role in promoting the re-growth and survival of cancer cells after chemotherapy treatment.

**Figure 5.**
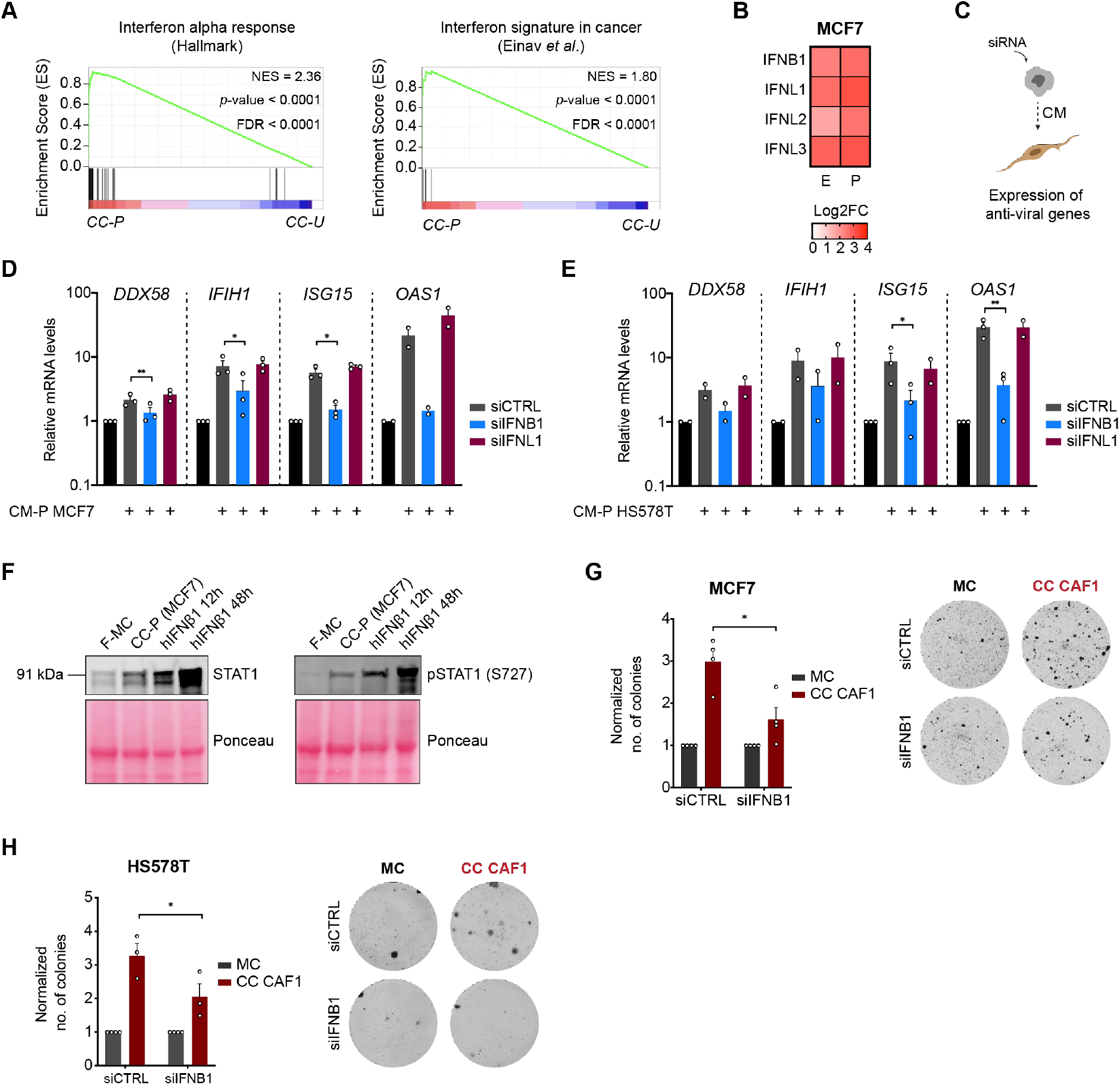
IFNβ1 secreted by chemotherapy-treated cancer cells drives fibroblasts into an anti-viral state and promotes recovery. **A**: GSEA of interferon response (HALLMARK) and interferon signature in cancer^55^ in fibroblasts in co-culture with paclitaxel-treated cancer cells (CC-E) compared with untreated cancer cells (CC-U). NES = normalized enrichment score, FDR = false discovery rate. P values were determined by random permutation tests. **B**: Heat-map with expression of significantly upregulated interferon transcripts in MCF7. Results for epirubicin- (E) and paclitaxel- (P) treated cancer cells are shown in log2 fold change and are normalised to untreated MCF7. **C**: Layout of experimental setup for interferon KD experiments. **D-E**: *DDX58, IFIH1, ISG15* and *OAS1* expression in CAF1 determined by RT-qPCR. CM from paclitaxel-treated MCF7 (**D**) and HS578T (**E**) transfected with a control siRNA (siCTRL), a siRNA against *IFNB1* (siIFNB1) or a siRNA against *IFNL1* (siIFNL1) was collected and added to CAF1. Relative fold change is normalised to CAF1 grown in DMEM (black bar). mRNA levels were normalized against two house-keeping genes (*ACTB* and *PUM1*). Each dot represents an independent experiment. P values for n>2 were calculated using unpaired two tailed t-test in biological replicates. * p<0.05, ** p<0.01. **F**: Immunoblot of STAT1 and pSTAT1 (S727) in CAF1 lysates in mono-culture (F-MC), in coculture with paclitaxel-treated MCF7 (CC-P) for 5 days or stimulated with 100 ng/mL of recombinant human IFNβ1 (hIFNβ1) for 12h or for 48h. Ponceau was used as loading control. **G-H**: Quantification of colonies in recovery assay of MCF7 (**G**) and HS578T (**H**) transfected with a siCTRL or a siRNA against *IFNB1* (siIFNB1). Representative pictures of wells at the end point are shown on the right panels. Each dot represents an independent replicate (n=4 for MCF7 and n=3 for HS578T). P value was calculated using paired two-tailed t-test in biological replicates for MCF7 (**G**) and one-tailed t-test in biological replicates for HS578T (**H**). * p<0.05. Data is shown as mean ± SEM.

Together we show that chemotherapeutic drugs induced the expression of IFNβ1 in cancer cells and that secreted IFNβ1 induced the expression of interferon stimulated genes in fibroblasts. Importantly, this crosstalk promoted the recovery of cancer cells after chemotherapy treatment.

### Clinical significance of IFNβ1 and anti-viral state

To explore the clinical significance of this newly discovered axis of communication between cancer cells and fibroblasts, publicly available datasets of patient data were investigated. To this end, we first tested the potential correlation between the levels of *IFNB1* and the outcome of patients using ROC plotter^31^. Patients that were treated with either taxanes or anthracyclines were selected for the analysis (Fig 6A). High levels of *IFNB1* were associated with a worse outcome in recurrence free survival (RFS) (Fig 6B-C), indicating an inverse correlation between RFS time and *IFNB1* expression. We next clustered breast cancer patients that had been treated in a neoadjuvant setting with a taxane and anthracycline-based regimen according to the expression of the antiviral signature identified in the fibroblasts – IFN signature (IFNS) (<u>Supplementary Table 2</u> for full list of genes). High expression of the IFN signature genes was associated with poor relapse-free survival also in this cohort (Fig 6D).

**Figure 6.**
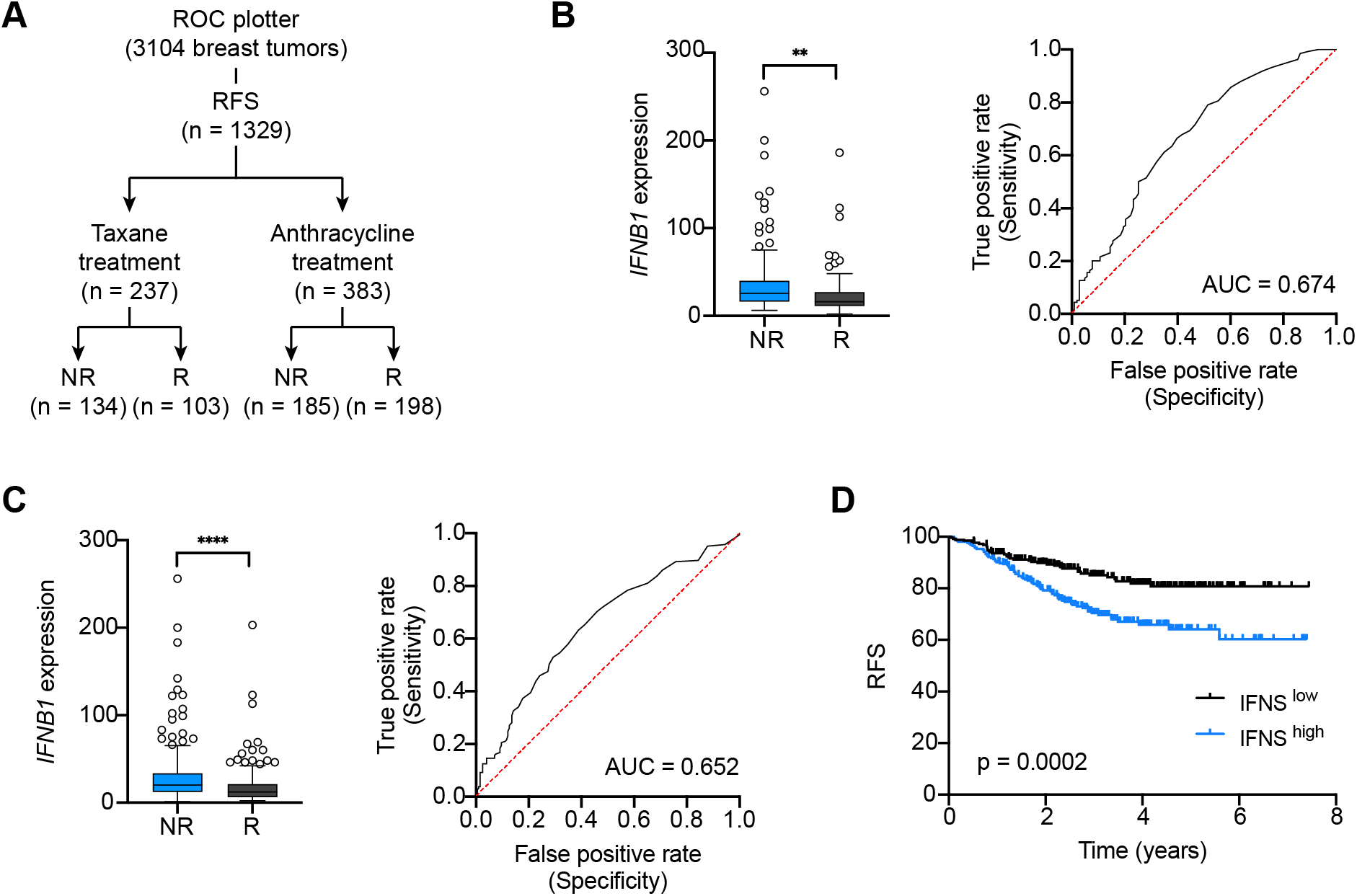
Clinical significance of IFNβ1 axis. **A**: Schematic overview of patient number for each group in ROC plotter. **B-C**: Analysis of *IFNB1* expression and RFS outcome using ROC plotter in breast cancer patients treated with taxanes (**B**) or anthracyclines (**C**). P value was calculated using unpaired two-tailed t-test. ** p<0.01, **** p<0.0001. NR = non-responder, R = responder. **D**: Kaplan-Meier analyses of breast cancer patients, associating IFN signature (IFNS) with recurrence free survival (Compiled data set from GSE25055 and GSE25065, n=508: IFNS^high^ n=258, IFNS^low^ n=250). Median cut-off was used to group samples into low and high. P values were determined by log-rank (Mantel-Cox) test.

We show that cancer cells exposed to chemotherapy upregulate the expression of IFNβ1 which is then secreted into the microenvironment and modulates the activation of fibroblasts. Fibroblasts exposed to IFNβ1 acquire an anti-viral state that is characterised by increased expression of numerous inflammatory modulators, which can then promote the recovery of cancer cells after chemotherapy (See Fig 7 for schematic overview).

**Figure 7.**
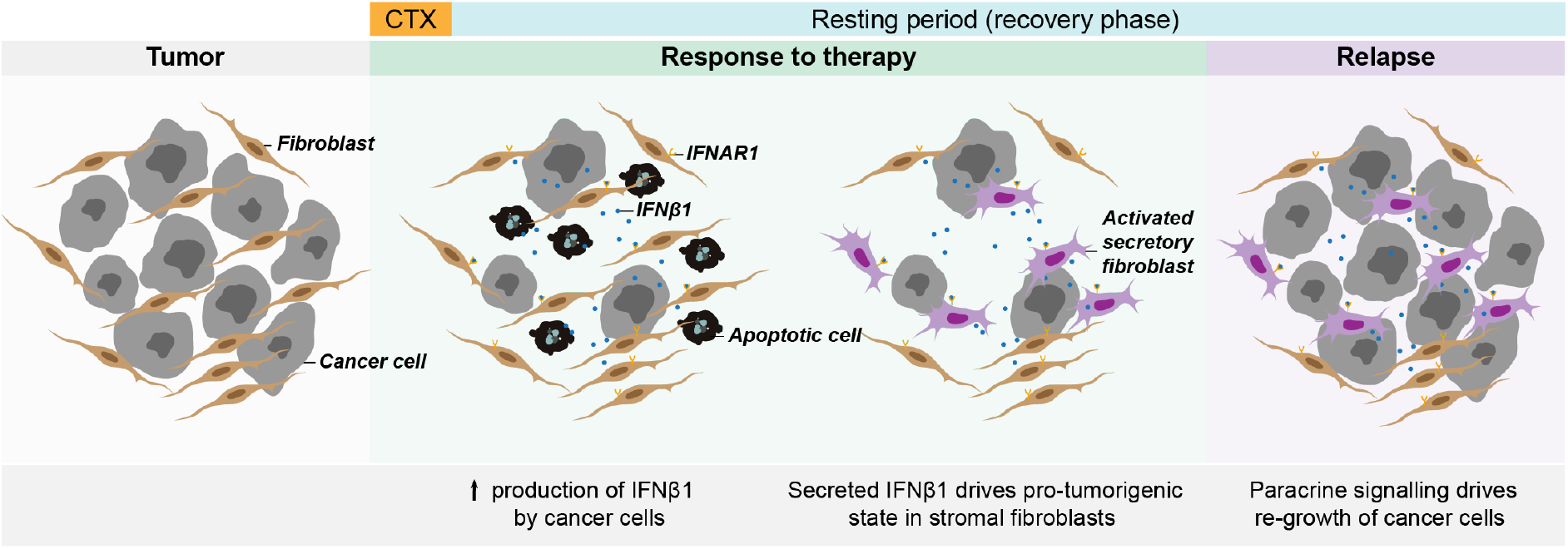
Schematic overview.

## Discussion

Chemotherapy is still the standard of care for a large number of patients with cancer. In breast cancer, a regimen based on anthracyclines and taxanes is commonly used^32^. Treatments are often given in cycles, in which patients are administered a maximum tolerable dose of the respective chemotherapeutic agent. The frequent failure of cancer cells to respond to therapy leads to tumour progression and ultimately results in the development of advanced metastatic disease, which is the main cause of death in cancer patients. An understanding of the mechanisms that drive therapeutic failure and development of resistance is of utmost importance and could help devising strategies to prevent relapses and metastatic disease. Cancer cells can evade therapy through a variety of mechanisms and the tumour microenvironment is known to play an important role in this process. In this study we have investigated the impact of high chemotherapeutic doses on cancer cells, its influence in the crosstalk with stromal fibroblasts and how this communication affects the outcome of cancer cells to chemotherapy treatment. Indeed, cancer-associated fibroblasts have been shown to drive the development of resistance of cancer cells to several therapies, including chemotherapy^5,6^. Here we show that stromal fibroblasts are not only affecting the response of tumour cells to chemotherapy by promoting the development of resistance, but they can also influence the fate of cancer cells after the exposure to cytotoxic agents. We provide evidence that in the presence of fibroblasts, cancer cells can recover from the cytotoxic damage induced by chemotherapy and re-start proliferating. This recovery potential is absent when cancer cells are grown alone. Additionally, we show that the cancer cells that recover in the presence of fibroblasts are not resistant to the agents that they were initially treated with. While the presence of resistant cells is one of the main drivers of therapeutic failure, it has been previously reported that persister cancer cells can escape the cytotoxic or cytostatic effect of treatment^3,33^. Based on the results from our study we hypothesize that fibroblasts help these cancer cells survive and re-enter cell cycle facilitating the occurrence of relapses.

In recent years, the secretome of cancer cells and its impact on therapy response has gained attention and numerous studies explore its role in several aspects of tumour progression. Molecules released by cancer cells can strongly modulate not only other cancer cells but also their microenvironment^7^. Several factors can influence and alter the spectrum of secreted molecules by cancer cells, including a large number of conventional cancer therapies. For example, it has been described that chemotherapy treatment can lead to the expression and release of several pro-tumourigenic molecules, including interleukins such as IL-6, whose expression negatively correlates with patient outcome^34,35^. The expression of other molecules, such as type I IFN, has also been shown to be modulated by anti-tumour therapies, especially radiation. Type I IFN signaling can have both anti-tumourigenic and pro-tumourigenic roles. Several studies demonstrate the importance of type I IFN signaling in effective anti-tumour immune responses^20,21,36^. Nevertheless, the role of type I IFN in the context of cancer is still not clear. While strong evidence exists that type I IFN is essential in several tumour entities for effective therapy response and immune engagement against cancer cells, other studies seem to challenge this black and white view on this signalling pathway. Chen *et al.* showed that depletion of type I IFN receptor, *IFNAR1,* in cancer cells promoted a strong immune response after treatment with radiation^12^, indicating that the outcome of type I IFN signalling in tumour progression might dependent on several factors, including in which compartment of the tumour this pathway is activated. Moreover, high expression of interferon-regulated genes in several tumour entities also strongly correlated with a worse outcome in patients^37,38^. All of these studies highlight the high degree of plasticity of type I IFN and the need for a better understanding of the role of this pathway in the context of cancer. While most of these studies focus on the impact of type I IFN in immune cells, much fewer reports addressing its effect in stromal fibroblasts are available. Of note, Hosein et al. identified in breast cancer a subset of fibroblasts characterized by a type I IFN response and showed that high expression of certain interferon stimulated genes in patient samples correlated with patient outcome^37^. In our study we show that cancer cells exposed to chemotherapeutic agents upregulate the expression of type I IFN, namely IFNβ1. This effect was observed when either a taxane (e.g. paclitaxel) or an anthracycline (e.g. epirubicin) were applied. Additionally, exposure of stromal primary fibroblasts to the conditioned-media of chemotherapy-treated cancer cells triggered the acquisition of an anti-viral, pro-inflammatory state in fibroblasts that was characterized by the expression of multiple ISGs. The expression of ISGs in the context of cancer has been previously shown to be induced by nucleic acid sensing pathways^5,22,39^. However, here we do not find a direct link between these pathways and the inflammatory response observed in fibroblasts, but instead we show that this state is mediated by cancer cell-secreted IFNβ1. Nonetheless, we speculate that the upregulation of IFNβ1 in cancer cells after treatment might be driven by the nucleic acid sensing pathway cGAS-cGAMP-STING, since accumulating evidence has shown that therapies that induce genomic instability, including radiation, can trigger the expression of interferons via this pathway^40,41^. This could also explain why chemotherapeutic agents with distinct mechanisms of action such as paclitaxel and epirubicin, induce the expression of IFNβ1. Despite the fact that these drugs are known to exert their cytotoxic effect via different ways, they both induce high levels of genomic instability. Moreover, we linked IFNβ1 to the increased recovery potential observed in cancer cells in the presence of fibroblasts. Silencing of *IFNβ1* in cancer cells abrogated their recovery capacity when they were cultured with stromal fibroblasts, indicating that IFNβ1 reprograms fibroblasts into a pro-tumourigenic state.

Inflammation is a complex process in cancer. Fibroblasts have been described as an important source of tumour-promoting inflammation with several studies finding evidences that pro-inflammatory fibroblasts drive malignancy and the development of metastsaes^29,42–44^. In our system, this anti-viral, pro-inflammatory state that is induced in fibroblasts by chemotherapy-treated cancer cells seems to promote the recovery of cancer cells after chemotherapy treatment. It was previously described that exposure of fibroblasts to high doses of chemotherapy could also drive an inflammatory microenvironment^28,45^. In contrast with these observations, in our system treatment of fibroblasts with the same chemotherapeutic doses that cancer cells were exposed to, did not drive the expression of ISGs in this cell type. In an attempt to more closely mimic the situation that is present in the tumour context, fibroblasts were grown to full confluency and the effect of chemotherapy in their cell viability was minimal. We thus propose that differences between the experimental setups used in our as compared to previous studies could explain the differences in the results obtained. Exposure of stromal fibroblasts to IFNβ1 led to the upregulation of specific cytokines, such as *CCL5* and *CXCL10.* The impact of IFNβ1 in the expression of these two cytokines is further supported by other studies^22,39^. *CXCL10* has been recently shown to promote a stemcell like phenotype in cancer cells^43^. In addition, another study has also connected pro-inflammatory cytokines with the expansion of stem-like cells and development of chemoresistance^28^, although they implicate different cytokines and chemokines in this process. Thus, we hypothesize that activated fibroblasts by IFNβ1 may drive the expansion of populations of stem-cell like cells that do not undergo apoptosis after the exposure to chemotherapy. It would be interesting to investigate if the cytokines that are secreted by IFNβ1-induced pro-inflammatory fibroblasts are driving the expansion of these populations. Moreover, these cytokines are known immune modulators and it would be important to further explore their impact on the immune milieu in this context.

We show that IFNβ1-induced pro-inflammatory fibroblasts promote the survival and recovery of cancer cells after chemotherapy treatment. Importantly, we find a negative correlation between IFNβ1 expression and RFS in breast cancer patients treated with taxane and anthracycline regimens thereby attaching clinical relevance to our findings. Furthermore, disease-free survival time in patients with high expression of the IFN signature identified in fibroblasts was significantly reduced, indicating that the activation of this signaling axis could be important in driving the appearance of relapses. Ultimately, this newly identified signaling axis should be further explored as its targeting might represent a potential strategy to improve the outcome of patients to chemotherapy treatment.

## Materials and Methods

### Isolation and establishment of primary culture of human fibroblasts

Generation of primary cultures from breast fibroblasts was done in collaboration with the National Centre for Tumour diseases (NCT) in Heidelberg and the Women’s Clinic in Tübingen. The scientific use of human tissue samples was approved by the medical faculty of the University Hospital Tübingen (ethical vote: 150/2018BO2) and by the Medical Faculty of Heidelberg (ethical vote: S-392/2015). Isolation of fibroblasts from fresh tissues was performed as previously reported^46^. Very briefly, tissue was digested in collagenase (Sigma-Aldrich, Missouri, USA) for two hours and the single cell suspension obtained was seeded in a cell culture dish. Preferential attachment to plastic by fibroblasts resulted in a population of cells enriched for this cell type. Five fibroblast lines were isolated and established for this study. Two pairs of fibroblasts isolated from the same patient from non-malignant (NF) and tumour sites (CAF) were obtained. Non-malignant fibroblasts were collected from a distal area of the tumour site within the same breast. Primary fibroblasts in experiments were used until they reached passage 8 or were in culture for a maximum of 1.5 months, after which they were discarded.

### Cell culture

All breast cancer cell lines were obtained from ATCC and regularly authenticated by multiplex cell line authentication (Multiplexion GmbH, Friedrichshafen, Germany) and tested for mycoplasma contamination by PCR. Breast cancer cell lines and primary human fibroblasts were grown in full growth media (DMEM supplemented with 10% FCS, 1% penicillin/streptavidin and 1% glutamine). MCF7-FUCCI cells were generated by infection with lentivirus-expressing mKO2-hCdt1(30/120)/pCSII-EF and mAG-hGeminin(1/110)/pCSII-EF. In all experiments, cells were kept in reduced serum media (DMEM supplemented with 2.5% FCS, 1% penicillin/streptavidin and 1% glutamine).

### Lentiviral infection

For generation of lentiviral particles, HEK293FT cells (Thermo Fischer Scientific, Germany) were co-transfected with either mKO2-hCdt1(30/120)/pCSII-EF or mAG-hGeminin(1/110)/pCSII-EF) and 2nd generation viral packaging plasmids VSV.G (Addgene #14888) and psPAX2 (Addgene #12260). 48 hours after transfection, virus containing supernatant was removed and cleared by centrifugation (5min/500g). The supernatant was passed through a 0.45 μm filter to remove remaining cellular debris. MCF7 cells were transduced with lentiviral particles at 75% confluency in the presence of 10 μg/ml polybrene (Merck, Germany). 24h after transduction virus containing medium was replaced with selection medium for the respective expression constructs to start the selection. After two weeks of selection, double positive cells for RFP and GFP were further enriched using FACS.

### Drug response

In a black 96-well plate (Greiner Bio-one, Kremsmünster, Austria), 1500 – 2000 cells per well were plated (day 0). At day 1 reduced serum medium with different concentrations of epirubicin (Biomol GmbH, Hamburg, Germany) and paclitaxel (Biomol GmbH, Hamburg, Germany) were added for three days. At defined time points cells were imaged and analysed using the ImageXpress Micro Confocal microscope (Molecular Devices, California, USA).

### Colony formation (recovery assays)

Cancer cells were seeded at a low density (5 x 10^4^ cells per well) in a 6-well companion plate (Corning, New York, USA). The next day, chemotherapeutic agents were diluted according to <u>Supplementary Table 3</u> in reduced serum media (DMEM supplemented with 2.5% FCS, 1% P/S and 1% glutamine) and added to the cancer cells for 3 days. 2 x 10^4^ primary fibroblasts were seeded in a 6.5 mm trans-well with 0.4 μm pore (Corning, New York, USA). After the three days incubation time, cancer cells were washed three times with PBS and allowed to recovery in reduced serum media in the absence (mono-culture) or presence of a trans-well with fibroblasts (co-culture). Media was refreshed every 4-5 days. After a minimum recovery period of 15 days, cancer cells were fixed in methanol for 10 min at room-temperature (RT) and stained with crystal violet for 1 hour. Crystal violet was removed, the wells were washed with distilled water and allowed to dry overnight. Plates were then scanned using an EPSON Perfection scan and images were analysed using Image J (NIH, USA).

### Conditioned-media

MCF7 (2 x 10^5^ cells per well) and HS578T (1.125 x 10^5^ cells per well) were seeded in 6-well plates and treated with 4 nM and 8 nM paclitaxel, respectively. After three days of incubation with chemotherapeutic drugs, cells were washed and fresh reduced serum media was added. 72 hours conditioned-media from cancer cells was collected and filtered with a 0.45 μm filter (Millex, Merck Millipore, Massachusetts, USA) before adding to fibroblasts. If additional treatment of conditioned-media was done, the supernatant was split into four conical tubes and incubated with either DNase I (Sigma-Aldrich, Missouri, USA), RNase A (Qiagen, Hilden, Germany), at 95°C or just kept at RT. DNase I and RNase A incubation was done for 30-45 min at RT and CM was kept at 95°C for no longer than 5 min. Afterwards, the pre-treated conditioned-media was added to fibroblasts that had previously been seeded in a 6-well plate at a confluency of 2 x 10^4^ cells. Fibroblasts were incubated for two days with conditioned-media and their cell pellets were collected for gene expression analysis. For exosome-depletion, cells were grown in DMEM supplemented with exosome-depleted FCS. 72 hours conditioned-media from cancer cells was collected and ultracentrifuged for 2 hours at 100,000x g. Supernatant was collected to a new tube and the exosome-enriched pellet fraction was resuspended and both fractions were added to the fibroblasts.

### Type-I interferon blocking antibodies

For blocking antibody treatment, supernatant from paclitaxel-treated cancer cells was collected and an anti-IFNB1 (R&D, AF814-SP) antibody at a concentration of 0.2 μg/mL was added. Fibroblasts were cultured in the supernatant of cancer cells for 48 hours before being collected for gene expression analyses.

### Immunofluorescence

For the characterization of primary fibroblasts isolated from patients, cells were seeded in glass coverslips. The following day, cells were fixed with 4% paraformaldehyde for 10 min at room temperature, permeabilized with 0.125% Triton X-100 for 10 min and blocked with 4% BSA for 1 hour. Primary antibodies against Fibronectin (ab2413, 1:100, Abcam) and Vimentin (sc-6260, 1:50, Santa Cruz), were incubated overnight in a humidified chamber at 4°C. Coverslips were washed three times with PBS containing 0.1% Tween-20 and incubated for 1 hour at room temperature with the respective secondary antibody (1:400, Abcam). Finally, coverslips were again washed and mounted in ProLong Diamong Antifade mounting media containing DAPI (Thermo Fisher Scientific, Massachusetts, USA). Samples were imaged with Zeiss Cell Observer inverted microscope and processed using Image J (NIH, USA).

### Flow cytometry

Analysis of expression of the activation marker aSMA was done in the fibroblast lines isolated from patient samples using flow cytometry. Primary fibroblasts were grown in reduced serum media for three days after which they were trypsinised and collected for antibody staining. Briefly, 1 x 10^4^ cells were washed one time with PBS and fixed with CytoFix/CytoPerm solution (BD Biosciences. USA) for 10 min on ice. Afterwards, cells were washed one time with 500 μL of 1x Perm/Wash Buffer (BS Biosciences, USA) diluted in PBS and incubated with 0.02 μg of aSMA (#50-9760-82, eBiosicence, Thermo Fisher Scientific, Massachusetts, USA) antibody for 30 min on ice. One more washing step with 1x Perm/Wash Buffer (BD Biosciences) was done and cells were acquired in the flow cytometer (BD FACS Canto II). Results were analysed using BD FACS DIVA (BD Biosciences, USA).

### Live cell imaging

5 x 10^4^ MCF7-FUCCI cells were seeded in a 6-well plate in full growth media. After cells attached to the well bottom, growth media was replaced by Leibovitz’s L-15 medium (Gibco, Thermo Fisher Scientific, Massachusetts, USA) supplemented with 2.5% FCS and cells were imaged every 20 min overnight with the motorised widefield microscope Cell Observer (Zeiss, Oberkochen, Germany). Images were analysed and processed using Image J (NIH, USA).

### Apoptosis analysis

Cancer cells were seeded (5 x 10^4^ cells per well), treated with either epirubicin or paclitaxel for three days and allowed to recover in the absence of drugs either in mono-culture or co-culture with fibroblasts. At selected time points, supernatant and attached cells were collected and stained with DAPI for 5 min before acquisition in BD FACS Canto II flow cytometer (BD Biosciences, USA). Results were analysed using the BD FACS DIVA (BD Biosciences, USA).

### Cell cycle

Investigation of the cell cycle profile of cancer cells was done using 5-ethynyl-2’-deoxyuridine (EdU) labelling and the FUCCI system. Cancer cells were seeded (5 x 10^4^ cells per well), treated with chemotherapy for three days and allowed to recover in the absence of drugs either in mono-culture or co-culture with fibroblasts. At selected time points, 10 μM of EdU was diluted in reduced serum media and added for 30 hours to cultures. After the incubation time, cancer cells were fixed in 4% PFA for 20 min at RT and stained according to the Click-iT EdU imaging kit protocol (Invitrogen, California, USA). Stained cancer cells were imaged and quantified using the ImageXpress Micro Confocal microscope (Molecular Devices, California, USA). FUCCI system analysis was done using flow cytometry. Briefly, MCF7-FUCCI cells were seeded and submitted to the same experimental setup described above. Cells were collected by trypsinisation and the channels RFP and GFP were acquired in BD FACS Canto II flow cytometer (BD Biosciences, USA). Results were analysed using the BD FACS DIVA (BD Biosciences, USA).

### Senescence

Chemotherapy-treated MCF7 in mono-culture and co-culture with fibroblasts were fixed at different time-points of recovery in 4% PFA for 10 min. Cells were stained for SA-β-galactosidase activity overnight using a Cellular Senescence Assay kit (Merck Millipore, Massachusetts, USA) and imaged the next day in the motorised widefield microscope Cell Observer (Zeiss, Oberkochen, Germany). Images were analysed and quantifying using Image J (NIH, USA).

### RNA sequencing

Total RNA was isolated from cells using the RNeasy Mini Kit (Qiagen, Hilden, Germany) according to manufacturer’s protocol. RNA concentration and purity were analysed using NanoDrop 2000 spectrophotometer (Thermo Fisher Scientific, Massachusetts, USA). Samples were sequenced in the DKFZ Genomics and Proteomics Core Facility using Illumina platform HiSeq 4000 v4 single-read 50bp. Reads were mapped to the human reference genome (build 37, version hs37d5) using STAR (version 2.5.2b)^47^ and a 2-pass alignment. Duplicate reads were marked with sambamba (version 0.6.5)^48^ using 8 threads. Then, reads were sorted by position using SAMtools (version 1.6)^49^. BAM file indexes were generated using sambamba. Quality control analysis was performed using the SAMtools flagstat command, and the rnaseqc tool (version v1.1.8.1) with the 1000 Genomes assembly and gencode 19 gene models. FeatureCounts (version 1.5.1)^50^ was used to perform gene specific read counting over exon features based on the gencode V19 gene model. The quality threshold was set to 255 (which indicates that STAR found a unique alignment). For total library abundance calculations, during FPKM/TPM expression values estimations, all genes on chromosomes X, Y, MT, and rRNA and tRNA were omitted, as they possibly introduce library size estimation biases.

### Transfections

Transfections were done using Lipofectamine RNAimax (Invitrogen, California, USA) according to manufacturer’s instructions. siRNAs (Dharmacon, Colorado, USA) were used at a final concentration of 20 nM and are shown in <u>Supplementary Table 4</u>. Briefly, the day after cells were seeded, a mix containing the transfection reagents and respective siRNAs was added dropwise to the cells. Cells were incubated for 6 hours with the transfection mix, after which the media was aspirated and fresh media was added.

### RNA extraction

Total RNA was isolated from cells using the RNeasy Mini Kit (Qiagen, Hilden, Germany) according to manufacturer’s protocol. RNA concentration and purity were analysed using NanoDrop 2000.

### Reverse transcription quantitative PCR (RT-qPCR)

cDNA synthesis was done using the RevertAid H Minus First Strand cDNA Sythesis Kit (Thermo Fisher Scientific, Massachusetts, USA). Primers for gene expression analysis were designed using the Roche UPL Design Center (Roche Applied Science, Penzberg, Germany) and are shown in <u>Supplementary Table 4</u>. All experiments were performed in triplicates and relative quantification was done using the 2^-ΔCt^ method. Relative expression was normalized to the house-keeping genes *ACTB* and *PUM1.*

### Western-Blot

Fibroblasts were harvested and stored at −80°C. Cell pellets were lysed in Lysis Buffer containing 20 mM Tris/HCl (pH7.5), 150 mM NaCl, 1% Triton X-100, 1 mM DTT, 2 mM EDTA, 1 mM NaF, 1 mM Ortho-Vanadate, 25 mM beta-glycerophosphate and protease inhibitor cocktail (Roche Applied Science, Penzberg, Germany). Protein concentration of the samples was measured with BCA protein assay kit (Thermo Fisher Scientific, Massachusetts, USA) and quantified with GloMax microplate reader (Promega GmbH, Walldorf, Germany). Samples were loaded in a polyacrylamide gel and separated by SDS-PAGE. Proteins were transferred from the polyacrylamide gel to a PVDF membrane (Merck Millipore, Massachusetts, USA). Membranes were blocked with Rockland blocking buffer (Biomol GmbH, Hamburg, Germany) for 1 hour at RT and incubated with the primary antibodies (<u>Supplementary Table 6</u>) overnight at 4°C. After washing a minimum of three times for at least 15 minutes with 0.1% Tween in TBS (TBS-T) the membrane was incubated with a fluorescently labelled-secondary antibody (Goat anti-Rabbit Alexa Fluor 680 #A32734 against STAT1; Goat-anti-Mouse Alexa Fluor 800 #A32730 against ACTB) (Thermo Fisher Scientific), Massachusetts, USA) at RT for 1 hour. Finally, the membranes were washed with TBS-T and scanned with Odyssey infrared imaging system (LI-COR Biosciences, Nebraska, USA). Images were then processed using Image J (NIH, USA).

### Enzyme-linked immunosorbent assay (ELISA)

ELISA was used to quantify the amounts of IFNβ1 in the supernatant of cancer cells to validate the IFNβ1 silencing by the interference RNA. MCF7 was seeded in full growth media in a 6-well plate at a confluency of 2 x 10^5^. The next day, siIFNβ1 plus transfection mix was added to the cells and incubated for 6 hours. Transfection media was aspirated and reduced serum media with 10 μg/mL of Poly(I:C) (InvivoGen, California, USA) was added to the cells for three days. Secreted IFNβ1 was detected with human IFN-beta DuoSet ELISA kit (R&D systems, Minnesota, USA) according to manufacturer’s instructions. Optical density was measured at 450 nm using GloMax microplate reader (Promega GmbH, Walldorf, Germany). A standard curve was generated using the absorbance values of the standards and used to calculate the concentration of IFNβ1 in each sample.

### Gene set enrichment analysis

Gene set enrichment analysis was carried out using GSEA v.4.0.3^51,52^. Comparison between groups (CAF1_CC-U vs. CAF1_CC-P; MCF7_MC-U vs. MCF7_MC-E/P) was done using gene sets extracted from GSEA. All parameters were kept as default. Nominal P values were calculated based on random genes permutations.

### ROC plotter analysis

Correlation between relapse free survival (RFS) at 5 years and *IFNB1* expression in chemotherapy-treated breast cancer patients was done using ROC plotter^31^. Patients were selected for taxane or anthracycline treatment and the relative expression levels of *IFNB1* (208173_at) investigated in patients pre-defined as responders (R) or nonresponders (NR). No additional filter was applied and no outliers were excluded from the analysis.

### Kaplan-Meier survival analysis

Survival analysis was done using compiled breast cancer datasets of patients treated with neoadjuvant chemotherapy (GSE25055 and GSE25065). Samples were divided into IFNS high and low based on median cutoff, using mean expression of the 52 IFN signature genes (shown in <u>Supplementary Table 6</u>).

### Statistics and reproducibility

Statistical analysis was performed using GraphPad Prism software v8. Graphs are shown as mean ± SEM and each dot shown represents an independent biological replicate. Statistical analysis was performed in experiments with n≥3. Statistical tests used in experiments are reported in figure legends. P values ≤0.05 were considered statistically significant and statistical tests were two-tailed. For Kaplan-Meier analyses of breast cancer patients, statistical differences in survival curves were calculated by log-rank (Mantel–Cox) test.

## Abbreviations

BUB1: Budding Uninhibited by Benzimidazoles 1
CAF: Cancer-associated fibroblast
CC: Co-culture
CCL5: C-C Motif Chemokine Ligand 5
CDT1: Chromatin Licensing and DNA Replication Factor 1
CTX: Chemotherapy
CXCL10: C-X-C Motif Chemokine Ligand 10
DAMPs: Damage-associated molecular patterns
DAPI: 4’,6-diamidino-2-phenylindole
DNA: Deoxyribonucleic acid
DDX58: DExD/H-Box Helicase 58
EdU: 5-Ethynyl-2’-deoxyuridine
ELISA: Enzyme-linked immunosorbent assay
FUCCI: Fluorescent Ubiquitination-based Cell Cycle Indicator
GSEA: Gene set enrichment analysis
GFP: Green fluorescent protein
GO: Gene ontology
IFIH1: Interferon Induced with Helicase C Domain 1
IFNA: Interferon alpha
IFNAR1: Interferon alpha receptor 1
IFNB1: Interferon beta 1
IFNG: Interferon gamma
IFNL: Interferon lambda
IFNS: Interferon signature
IL6: Interleukin 6
IL10RA: Interleukin 10 receptor subunit alpha
IL28RB: Interleukin 28 receptor subunit beta
IRF: Interferon regulatory factors
IRSE: Interferon responsive sequence element
ISG15: Interferon-Stimulated Protein, 15 KDa
ISG: Interferon-stimulated genes
IFN: Interferon
KD: Knock-down
MC: Mono-culture
NF: Normal fibroblast
OAS1: 2’-5’-Oligoadenylate Synthetase 1
PAMPs: Pathogen-associated molecular patterns
PRRs: Pattern recognition receptors
RFS: Recurrence free survival
RFP: Red fluorescent protein
RNA: Ribonucleic acid
RT-qPCR: Reverse transcription quantitative PCR
siRNA: short interference RNA
STAT: Signal transducer and activator of transcription
TTK: Threonine tyrosine kinase

## Acknowledgements

We thank C. Körner for critical reading of the manuscript. Additionally, we want to thank D. Heiss for her technical assistance with the Western-Blots. We would also like to thank I. Zörnig and P. Schrotz-King from the Jäger lab for providing tissue and M. Berdiel-Acer for the isolation of primary fibroblasts. We are grateful to the Genomics and Proteomics, Omics IT and Data Management Core Facility as well as the Light Microscopy and the Flow Cytometry Core Facilities at the German Cancer Research Center for providing excellent services and technical support. A.M. was supported by a scholarship from the Helmholtz International Graduate School for Cancer Research.

## Author Contributions

A.M. designed and performed the experimental work in the study. Z.G. performed the bioinformatic analysis of RNA-sequencing data under the supervision of M.S. Gene set enrichment analyses and patient data set analyses was done by A.M. A.K. obtained breast cancer patient samples, performed isolation of primary fibroblasts and contributed with critical feedback. R.W. generated the MCF7-FUCCI cell line. A.M. and S.W. designed the study and wrote the manuscript. All authors read and approved the final manuscript.

## Supplementary Figures

**Supplementary Figure 1.**
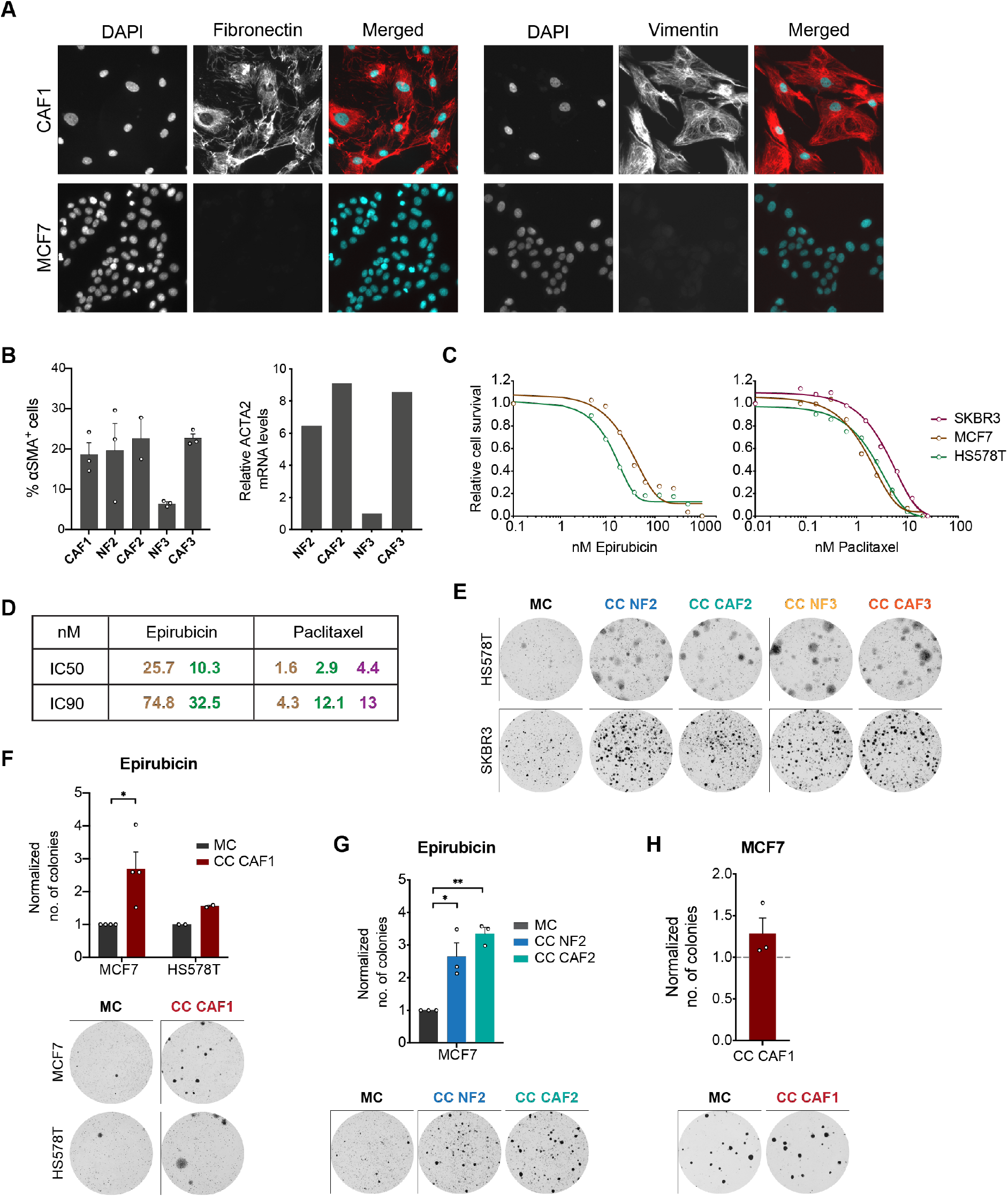
Fibroblasts promote the recovery of cancer cells after high-dose chemotherapy. **A**: Representative images of Fibronectin and Vimentin expression in primary fibroblasts (CAF1) and in MCF7 determined by immunofluorescence. **B**: Quantification of aSMA expression in the different fibroblast lines. Left panel shows results from flow cytometry analysis. Positive cells were determined based on isotype control levels. Each dot represents an independent replicate (For all fibroblasts, except CAF2: n=3; CAF2: n=2). Right panel shows ACTA2 mRNA levels in the several fibroblasts determined by RT-qPCR (n=1). **C-D**: Drug response curves of MCF7, HS578T and SKBR3 to epirubicin (left) and paclitaxel (right) determined by nuclei count. Results from three biological replicates (n=3) are shown as mean without variation for visual clarity (**A**). IC50 and IC90 values were calculated in GraphPad Prism 8.0 for the different cell lines and are shown in **B**. **E:** Representative images of the colony formation assay of HS578T and SKBR3 exposed to 8 nM paclitaxel in MC and CC with fibroblast pair #2 and #3 (n=1). **F**: Quantification of MCF7 and HS578T colonies in MC and CC with CAF1 at the end of recovery after treatment with epirubicin. Each dot represents an independent replicate (MCF7: n=6; HS578T: n=2). Representative pictures of colony formation assay wells at end-point are shown bellow. **G**: Recovery assay of MCF7 exposed to 70 nM epirubicin in MC or CC with CAF2 or NF2. Quantification of the number of colonies after 15 days of recovery for three independent experiments (n=3) is shown in top panel and representative pictures in lower panel. **H**: Colony formation assay of untreated MCF7 in CC with CAF1. Quantification is normalized to number of colonies of untreated MCF7 in MC. Representative images from three independent experiments (n=3) are shown on the lower panel.

**Supplementary Figure 2.**
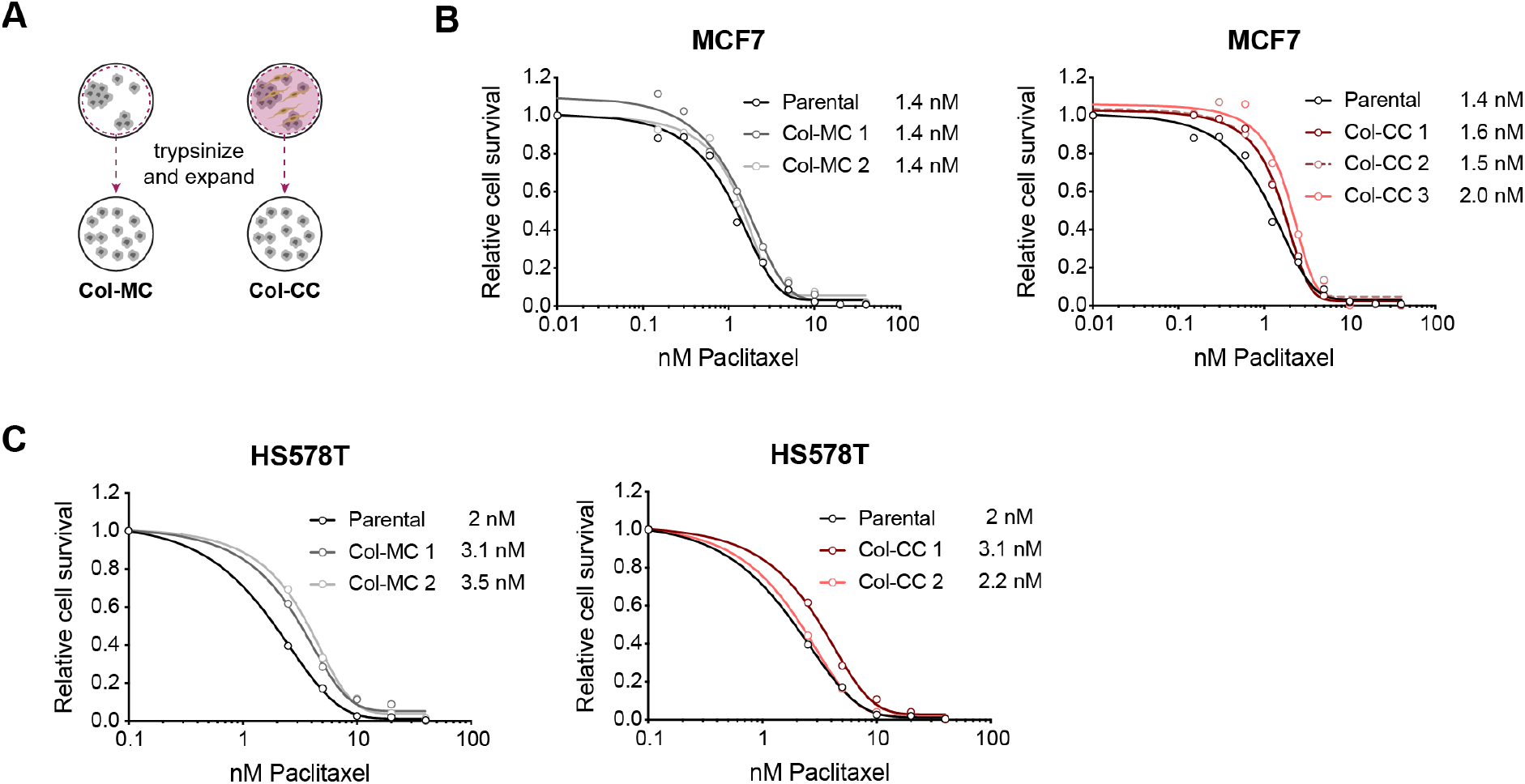
Cancer cells that recover are not resistant to chemotherapy. **A**: Schematic layout of colony analysis. Colonies from MC (Col-MC) and CC (Col-CC) were collected and expanded for drug response assay. **B-C**: Drug response curves to paclitaxel treatment of MCF7 (**B**) and HS578T (**C**) colonies determined by nuclei count. Results are shown as mean without variation for visual clarity. For MCF7, colonies from two and three independent experiments were collected for Col-MC (n=2) and Col-CC (n=3), respectively. For HS578T, colonies from two experiments for each condition were collected and used for the analysis (n=2). IC50 values were calculated in GraphPad Prism 8.0 for both cell lines.

**Supplementary Figure 3.**
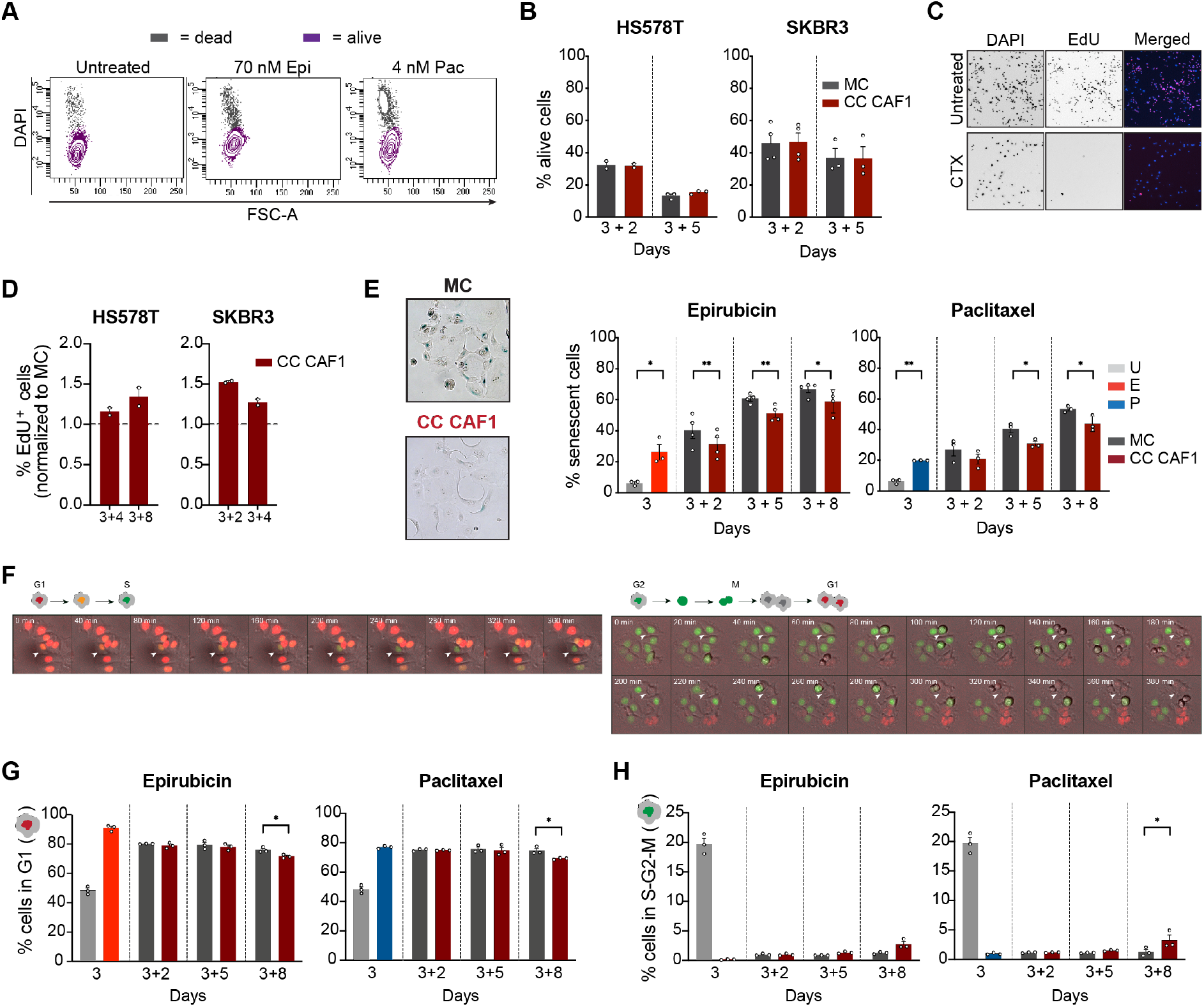
Fibroblasts promote the re-entering in cell cycle of cancer cells after chemotherapy treatment. **A-B**: Flow cytometry analysis of apoptosis levels in cancer cells. DAPI negative cells were quantified using flow cytometry, as seen in the representative plot shown in **A**. Quantification of DAPI negative cells in HS578T and SKBR3 for both MC and CC conditions after two (3+2) and five (3+5) days of recovery was done. Graphs in **B** show the percentage of alive cells (%) calculated by dividing the number of DAPI negative cells by the total number of singlets. Each dot represents an independent biological replicate. **C**: Representative pictures of the EdU labelling assay showing DAPI and EdU staining in untreated and chemotherapy-treated (CTX) MCF7. Ratio between the number of positive cells for EdU (EdU^+^) and total number of cells counted by DAPI staining was done to calculate the % of EdU^+^ cells after the cytotoxic stimuli with chemotherapy. **D**: Graphs show percentage of EdU positive cells in HS578T and SKBR3 in mono-culture versus co-culture with CAF1 at different time points of recovery after treatment with 8 nM paclitaxel (n=2 for each time point). **E**: SA-β-Galactosidase staining of MCF7. Left panel shows a representative picture of cells in MC and CC with fibroblasts. Quantification of the % of senescent cells from n≥3 is shown in the right panel. Data is shown as mean ± SEM and each dot represents an independent biological replicate. P values were calculated using unpaired two-tailed t-test on biological replicates. * p<0.05, ** p<0.01. **F**: Images from a time-lapse experiment of MCF7-FUCCI showing the different cell cycle phases transitions. White arrow shows transition from G1 to S phase (left panel) and G2-M-G1 transitions (right panel). Time stamp is shown in upper left corner of each picture. **G-H**: Quantification of Cdt1-RFP (G) and Geminin-GFP (H) expressing cells in MCF7-FUCCI by flow cytometry after treatment with epirubicin or paclitaxel at several recovery points. Same condition legend as shown in **E**. Data is shown as mean ± SEM from three independent experiments (n=3). P values were calculated using unpaired two-tailed t-test on biological replicates. * p<0.05.

**Supplementary Figure 4.**
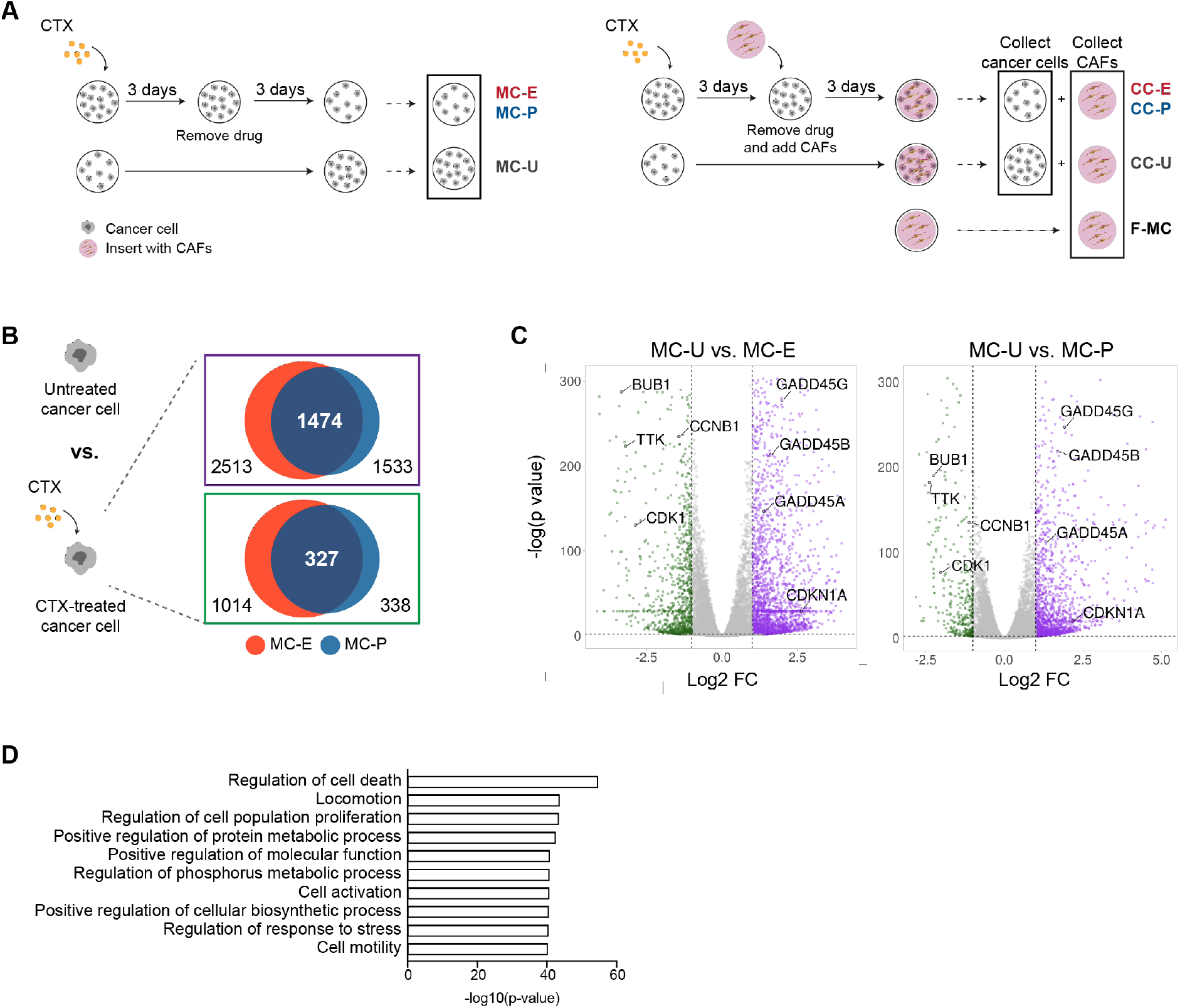
MCF7 and CAF1 RNA-sequencing. **A**: RNA-sequencing schematic layout. MCF7 treated with either epirubicin (E) or paclitaxel (P) were allowed to recover for three days in the absence (MC) or presence of fibroblasts (CC). At time point day 3+3, both MCF7 and CAF1 were collected and their gene expression profiles assessed by RNA-sequencing. Three replicates (n=3) per condition were sequenced. **B**: Venn diagram depicting differently upregulated (FC>2) (purple box) and downregulated (FC<0.5) (green box) genes in MCF7 after chemotherapy (CTX) treatment compared to untreated cells. Red circle represents the differently expressed genes after epirubicin treatment (MC-E) while the blue circle shows the ones in paclitaxel treatment (MC-P). **C**: Volcano plots showing the significantly differently expressed genes in CTX-treated MCF7 compared to untreated (MC-U). Upregulated genes are shown in purple and downregulated in green. Example of genes related to the p53 pathway (upregulated) and cell cycle progression (downregulated) are highlighted. Volcano plots were designed using the *VolcaNoseR^60^* tool. **D**: Analysis of data generated by RNA-sequencing using GSEA (GO: Biological process). Main pathways (top 10) upregulated in untreated MCF7 cells in co-culture with CAF1 compared to mono-culture are shown.

**Supplementary Figure 5.**
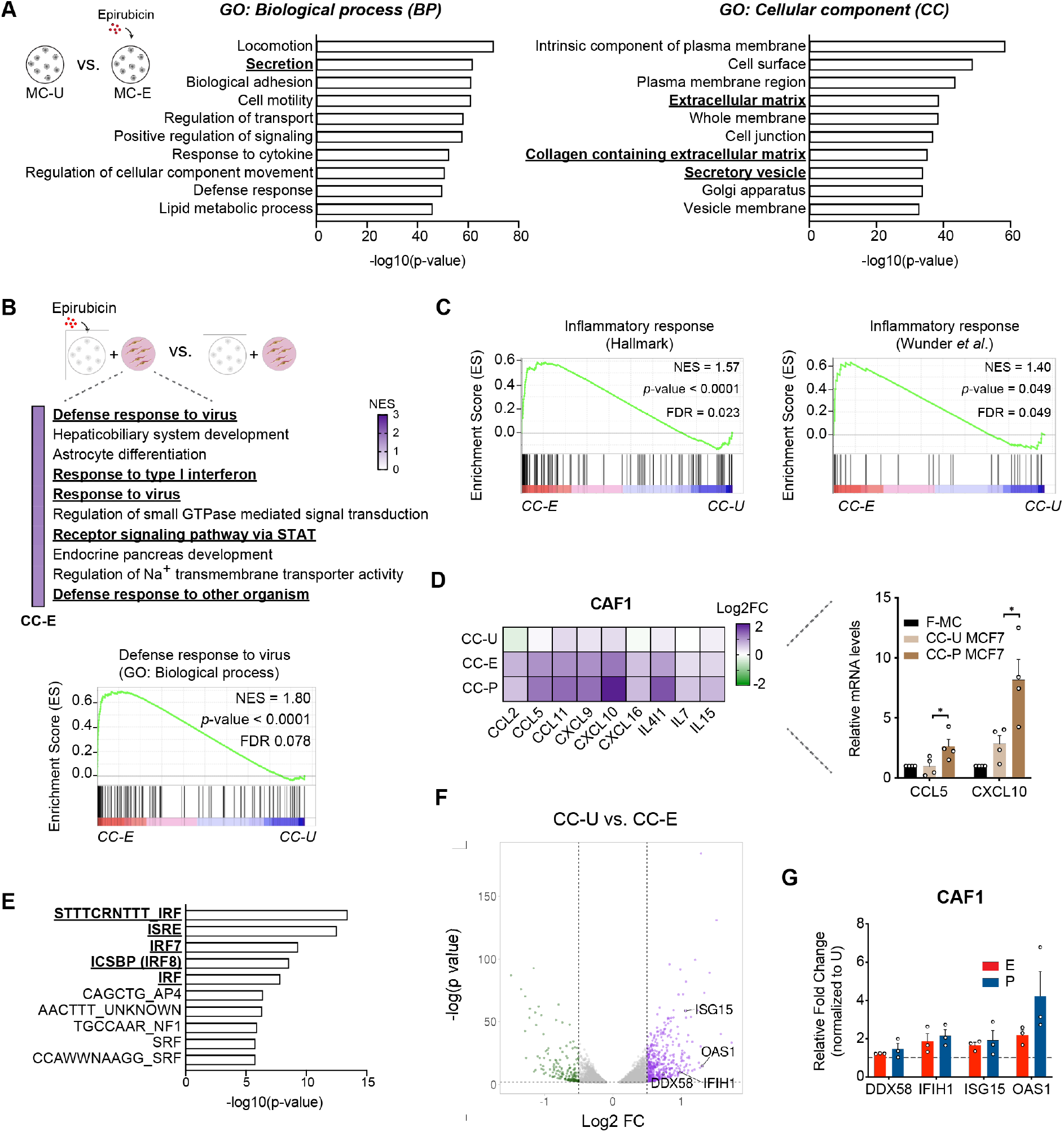
Fibroblasts acquire an anti-viral state after co-culture with chemotherapy-treated cancer cells. **A**: Top 10 enriched GO terms for biological process (BP) (left panel) and cellular component (CC) (right panel) of top 2000 upregulated genes in epirubicin-treated MCF7 compared to untreated. Terms related to secreted factors are highlighted in bold. **B**: GSEA analysis of upregulated genes in CAF1 in CC-E compared to CC-U. Top 10 significantly enriched GO terms enriched are shown. NES = normalized enriched score. Terms related to anti-viral response are highlighted. **C**: Enrichment for viral responses signatures in CAF1 in CC-E versus CC-U. FDR = false discovery rate. P values were calculated by random permutation tests. **D**: Heat-map with the expression of several pro-inflammatory cytokines in CAF1 in co-culture with untreated MCF7 (CC-U) and with epirubicin (CC-E) or paclitaxel-treated MCF7 (CC-P). Relative expression is normalised to CAF1 in mono-culture and is shown as log2 fold change. Right panel shows qRT-PCR analysis of the expression of *CCL5* and *CXCL10* in CAF1 grown for 5 days in mono-culture (F-MC) or co-culture with untreated (CC-U) or paclitaxel-treated (CC-P) MCF7. Values were normalized to CAF1 in F-MC. mRNA levels were normalized against two house-keeping genes *(ACTB* and *PUM1).* Each dot represents an independent experiment (n=4). P values were calculated using unpaired two-tailed t-test in biological replicates. * p<0.05. **E**: Top 10 over-represented transcription factors and motifs involved in the regulation of genes upregulated in CAF1 in CC-E. Transcription factors known to be involved in anti-viral response are underlined and highlighted in bold. **F**: Volcano plots of differently expressed genes in CAF1 in CC-E compared to CC-U. Upregulated genes are shown in purple and downregulated genes in green. Anti-viral genes *DDX58*, *IFIH1*, *ISG15* and *OAS1* are highlighted. Volcano plot was designed using *VolcaNoseR* tool. **G**: Anti-viral genes expression in CAF1 after treatment with 70 nM epirubicin (E) or 4 nM paclitaxel (P). Expression level was analysed using qRT-PCR and values normalized to untreated (U) CAF1. Data is shown as mean ± SEM from three independent experiments (n=3).

**Supplementary Figure 6.**
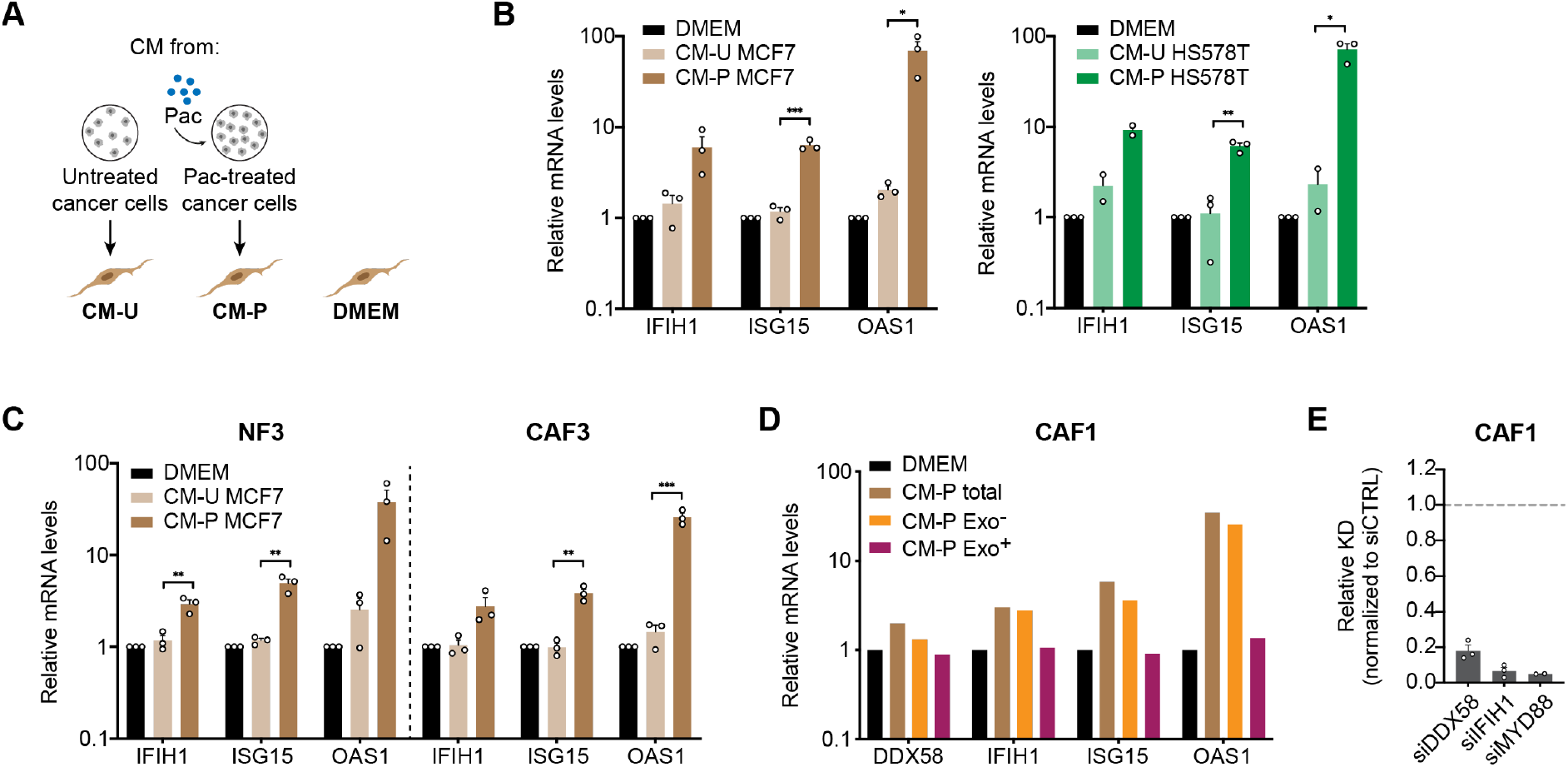
Anti-viral like state in fibroblasts is independent of nuclei acid sensing pathways. **A**: Schematic layout for conditioned-media experiment. Supernatant from untreated (CM-U) or paclitaxel-treated (CM-P) cancer cells was collected and added to primary fibroblasts. Fibroblasts grown in media are referred to as DMEM. **B**: Impact of MCF7 (left) and HS578T (right) conditioned-media in the expression of *IFIH1, ISG15* and *OAS1* in CAF1 determined by qRT-PCR. **C**: qRT-PCR analysis of *IFIH1, ISG15* and *OAS1* expression in fibroblast pair #3 (NF3 and CAF3). In **B** and **C** values were normalized to CAF1 grown in DMEM. mRNA levels were normalized against two house-keeping genes *(ACTB* and *PUM1).* Data is shown as mean ± SEM from at least two independent experiments (n≥2). Each dot represents a biological replicate. P values were calculated using unpaired two-tailed t-test. * p<0.05, ** p<0.01, *** p<0.001. **D**: Impact of exosomes in the anti-viral state of CAF1. Conditioned-media from paclitaxel-treated MCF7 was added directly to CAF1 (CM-P total) or ultra-centrifuged before. Supernatant fraction – exosome depleted (CM-P Exo^-^) – and pellet fraction – exosome enriched (CM-P Exo^+^) – were separated and added to CAF1. Expression of *DDX58, IFIH1, ISG15* and *OAS1* was assessed using RT-qPCR. Values were normalized to CAF1 grown in DMEM. mRNA levels were normalized against two house-keeping genes *(ACTB* and *PUM1*). Data from one experiment (n=1) is shown. **E**: Knock-down efficiency determined qRT-qPCR of the RIG-I pathway cytosolic receptors – *DDX58* and *IFIH1* – and of the TOLL-like receptor adaptor protein – *MYD88.* Values were normalized to expression in cells transfected with a control siRNA (siCTRL).

**Supplementary Figure 7.**
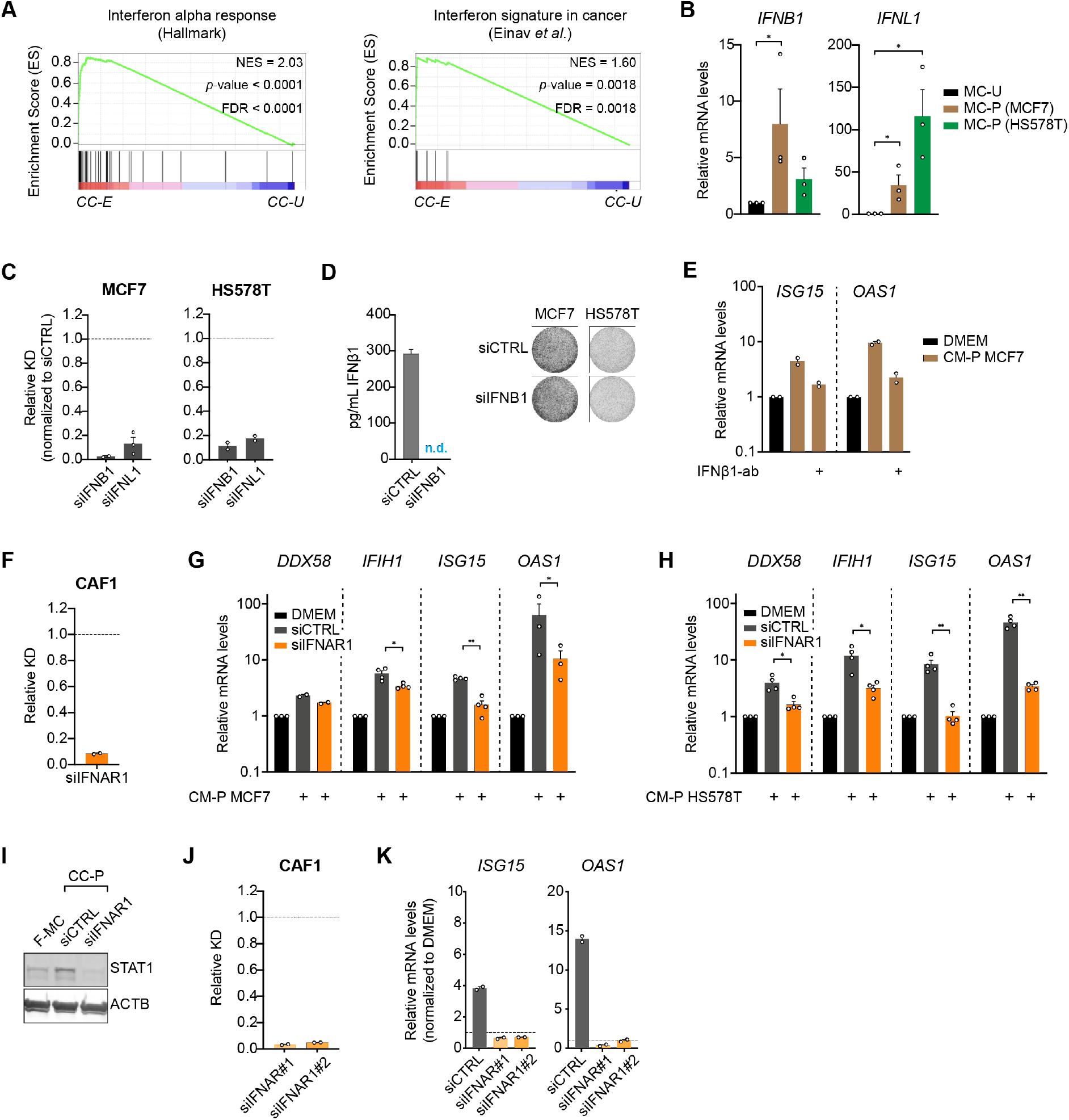
IFNβ1 secreted by chemotherapy-treated cancer cells drives fibroblasts into an anti-viral state. **A**: GSEA of interferon response (HALLMARK) and interferon signature^61^ in fibroblasts in coculture with epirubicin-treated cancer cells (CC-E) compared with untreated cancer cells (CC-U). NES = normalized enrichment score, FDR = false discovery rate. P values were determined by random permutation tests. **B**: qRT-PCR analysis of the expression of *IFNB1* and *IFNL1* in MCF7 and HS578T treated with paclitaxel (MC-P). Values were normalized to untreated cancer cells (MC-U). mRNA levels were normalized against two house-keeping genes *(ACTB* and *PUM1).* Each dot represents an independent experiment (n=3). P values were calculated using unpaired two-tailed t-test in biological replicates. * p<0.05. **C**: KD efficiency of siIFNB1 and siIFNL1 in MCF7 and HS578T, measured by qRT-PCR **D**: KD efficiency of siIFNB1 in MCF7, measured by ELISA. Gene expression values were normalized to cells transfected with a control siRNA (siCTRL). Pictures show representative wells of MCF7 and HS578T transfected either with siCTRL or siIFNB1 after six days of transfection. **E**: Impact of neutralizing antibody against IFNβ1 (IFNβ1-ab) in anti-viral gene expression in CAF1 measured by qRT-PCR. *ISG15* and *OAS1* expression in CM-P conditions were normalized to DMEM. Data from two independent experiments (n=2) is shown. **F**: KD efficiency of siIFNAR1 (pool of 4 independent siRNAs against IFNAR1) measured by qRT-PCR. Values were normalized to cells transfected with a control siRNA (siCTRL). **G-H**: qRT-PCR of anti-viral genes – *DDX58, IFIH1, ISG15* and *OAS1* – in CAF1 transfected with an siCTRL or siIFNAR1 (pool) and exposed to the supernatant of paclitaxel-treated MCF7 (**G**) and HS578T (**H**). Values were normalized to CAF1 grown in media (DMEM) which is shown in the graphs by the black bar. Each dot represents an independent biological replicate. **I**: Immunoblot of STAT1 in CAF1 in mono-culture (F-MC) or in co-culture with paclitaxel-treated MCF7 (CC-P) and transfected with a control siRNA (siCTRL) or the siIFNAR1 (pool). ACTB was used as a loading control. **J-K**: Two independent siRNAs against siIFNAR1 were used. KD efficiency and impact in *ISG15* and *OAS1* measured by qRT-PCR are shown in **J** and **K**, respectively. Data is from two independent experiments (n=2). In all qRT-PCR experiments, mRNA levels were normalized against two house-keeping genes *(ACTB* and *PUM1).* Data is shown as mean ± SEM. Each dot represents an independent replicate. P values were calculated for experiments with n>2 using unpaired two-tailed t-test. * p<0.05, ** p<0.01.

## Supplementary Tables legends

**Supplementary Table 1.** List of top 10 HALLMARKS and REACTOME terms in untreated, epirubicin- and paclitaxel-treated MCF7.

**Supplementary Table 2.** Anti-viral (IFN) signature genes.

**Supplementary Table 3**. Chemotherapy concentrations.

**Supplementary Table 4.** List of siRNAs.

**Supplementary Table 5.** RT-qPCR primers and probes.

**Supplementary Table 6.** Antibodies list.

